# Steroid hormones produced by coral-associated *Endozoicomonas* prime the coral immune response during thermal stress

**DOI:** 10.1101/2023.09.19.558257

**Authors:** Amin R. Mohamed, Michael A. Ochsenkühn, Trent D. Haydon, Salah Abdelrazig, Lisa S.Y. Coe, Gabriel Mochales-Riaño, Woonkee S. Jo, Alan R. Healy, David Abrego, Jean-Baptiste Raina, John A. Burt, Shady A. Amin

## Abstract

The coral microbiome is believed to play a critical role in sustaining corals and enabling adaptation to thermal stress. A ubiquitous group of coral-associated bacteria, known as *Endozoicomonas*, are hypothesized to provide essential metabolites to corals, though the exact roles these bacteria play during ambient or thermal stress conditions are largely unknown. We demonstrate that while *Endozoicomonas* synthesizes the iron-binding siderophore aerobactin *in vitro*, this trait may be suppressed *in hospite*. Integrated metagenomics, metabolomics, and lab validations reveal that within the coral holobiont *Endozoicomonas* is the only microbe capable of metabolizing coral-derived testosterone and converting ubiquitous cholesterol into testosterone and progesterone, important steroids that accumulate within the coral holobiont during thermal stress. Provision of testosterone and progesterone to corals during thermal stress induced transcriptional reprogramming of the coral innate immune response, promoting cellular survival and suggesting active microbial involvement in immune priming and stress resilience. These findings highlight a previously unrecognized mechanism of microbial contribution to coral resilience. Lastly, the capacity of bacteria to synthesize eukaryotic steroids implies deep evolutionary connections between microbial metabolism and the origin of steroid-mediated processes in early eukaryogenesis.

## Introduction

Corals form intricate associations with a diverse microbiome, collectively referred to as the coral holobiont^1^. This microbiome has been recognized as an essential contributor to coral health and resilience, especially during thermal stress^2–4^. While the symbiotic interactions between corals and their primary algal endosymbiont, Symbiodiniaceae, have been extensively studied^5–7^, the functional roles of bacteria in the coral holobiont remain largely unexplored^2,8^, despite their importance for coral health^3,9,10^. Among the myriad of bacteria associated with corals, the genus *Endozoicomonas* has received considerable attention due to its ubiquitous distribution across coral species and geographic locations^11–13^ and the correlation between its abundance and coral health^11,14,15^.

While *Endozoicomonas* are sometimes present in the coral mucus^16,17^, they are primarily localized within coral tissues as cell-associated microbial aggregates (CAMAs)^18^. Based on genomic predictions, *Endozoicomonas* spp. are hypothesized to play a crucial role in coral holobiont metabolism through the recycling of essential nutrients and biosynthesis of B-vitamins, amino acids and organosulfur compounds^11,19–21^. Indeed, when exposed to coral tissue extracts, *E. marisrubri* activates vitamins B_1_ and B_6_ biosynthesis and glycolytic processes^22^. However, despite recent efforts^13,21,23–25^, a definitive role for *Endozoicomonas* within the coral holobiont and the precise function underlying its symbiotic association is yet to be identified.

In this study, we combine multiomics data (16S rRNA gene amplicons, shotgun metagenomics, and untargeted metabolomics) to elucidate the functional roles of *Endozoicomonas* during coral response to extreme thermal conditions in the Arabian Gulf, the hottest sea on Earth with surface temperature exceeding 36°C^26^. To investigate symbiotic interactions in the coral holobiont, our sampling scheme included both healthy corals, representing stable host-microbiome associations, and corals infected with white syndrome, representing a dysbiotic state. Diseased samples, characterized by a marked reduction in the relative abundance of *Endozoicomonas*, provided a natural contrast to healthy corals that enabled us to determine the functional contribution of this taxon. By integrating metagenomic and metabolomic data, we uncover both shared and niche-specific metabolic pathways among members of the coral holobiont. Given the prominence of *Endozoicomonas* as a key member of the healthy coral microbiome, we leverage pangenomic analysis to delineate conserved and unique functions in *Endozoicomonas* lineages. We then perform additional experimental validations of these functional predictions.

We show that *Endozoicomonas* encodes genes for the biosynthesis and uptake of the iron-binding siderophore aerobactin, and confirm its ability to produce it *in vitro*, likely to access bioavailable iron *ex hospite*. In addition, we show that during thermal stress, two terminal steroid hormones, testosterone and progesterone, are abundant in corals despite the depletion of most of their precursors. We reveal that *Endozoicomonas* is the only holobiont member to possess the gene repertoire required for testosterone degradation and, surprisingly, can metabolize cholesterol to produce progesterone and testosterone *in vitro*, a unique capability only present in select bacteria. Finally, we show that progesterone and testosterone play a critical role in modulating the coral immune response during thermal stress by promoting pro-survival signaling while suppressing pro-inflammatory responses. These findings constitute a major breakthrough in our understanding of bacterial contributions to coral resilience and host–microbe interactions.

## Results

We monitored and sampled colonies of the coral *Acropora pharaonis*^27^ for 16S rRNA gene amplicons, shotgun metagenomics and untargeted metabolomics from Saadiyat reef on the Arabian Gulf off the coast of Abu Dhabi. Sampling occurred across a temperature gradient starting with early thermal stress showing preliminary signs of white syndrome disease on some colonies at 32°C (June), severe thermal stress and further white syndrome progression at 34°C (August), and recovery at 27°C (October) (Extended Data Table 1). Samples were collected from healthy coral tissues at all three temperatures, and from dysbiotic controls (i.e., white-syndrome lesion sites (L) and healthy appearing tissue adjacent to the lesion (AL)) at 34°C (Fig. 1a). All samples were extracted for metabolites and genomic DNA to profile the coral holobiont metabolome and microbiome composition and functional potential.

**Figure 1.**
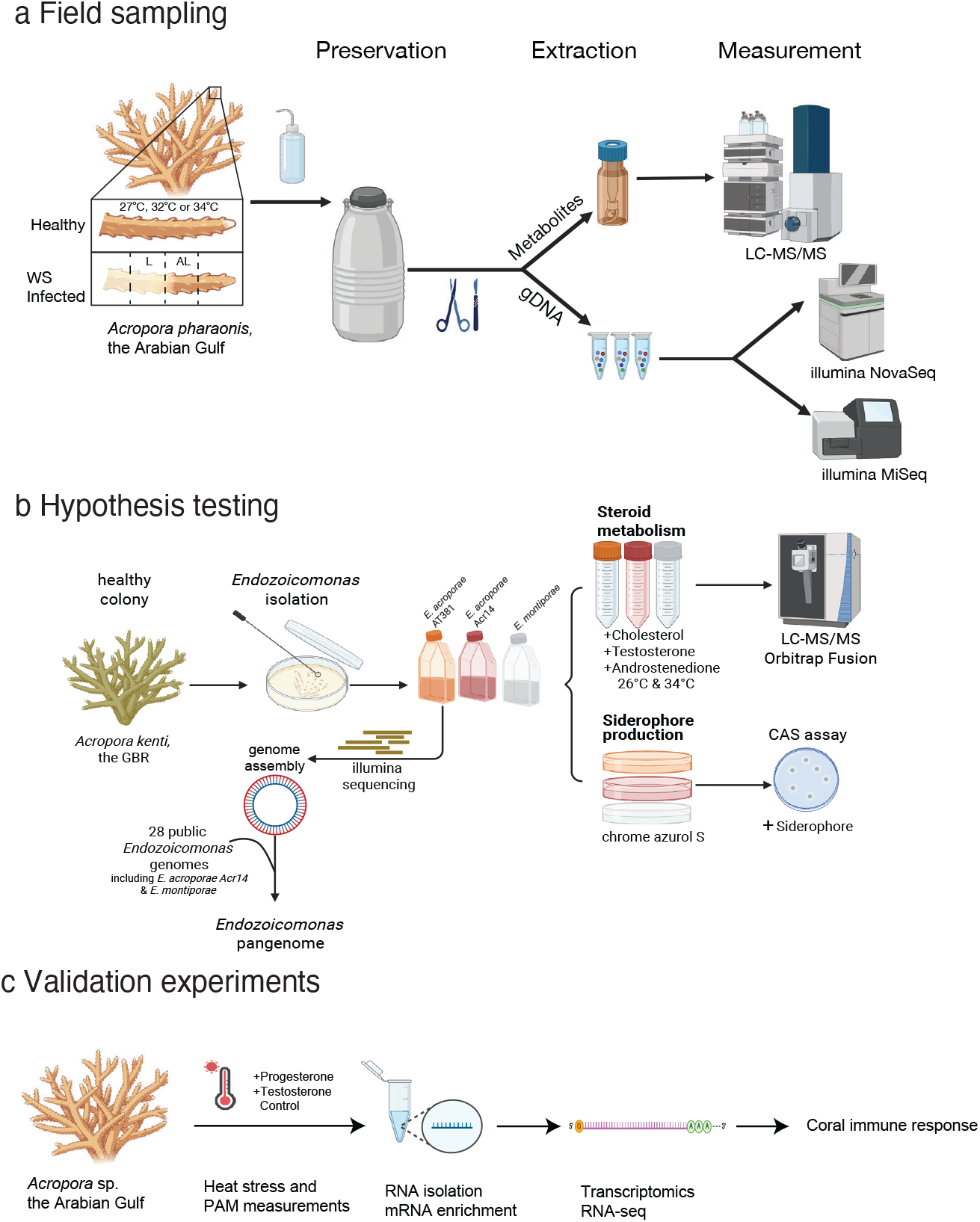
Sampling, analysis, and validation experiments schemes. **a**, *Acropora pharaonis* colonies were sampled across a temperature gradient before gDNA and metabolites were extracted for microbiome and metabolome analyses. Samples were collected during thermal stress at 32°C and 34°C (June/August) and recovery at 27°C (October). Additional samples showing white syndrome disease phenotypes at 34°C were also collected (L: lesion, AL: adjacent to the lesion). Samples were processed for 16S rRNA gene amplicon, shotgun metagenome, and untargeted metabolome analyses. **b**, To test the hypothesis derived from the multiomics data, experiments were conducted on three coral coral-associated *Endozoicomonas* to examine siderophore-mediated iron acquisition and steroid production. **c**, Transcriptomics was then used to examine the effect of steroid hormones on the coral immune response during thermal stress.

### Divergent metabolite and bacterial profiles in corals under extreme temperature

The intracellular metabolites of the coral holobiont were extracted and analyzed with reverse phase liquid chromatography-high resolution mass spectrometry (RP-LC/HRMS) in positive and negative ionization modes and in tandem with MS^2^ mass fragmentation. A total of 5878 putative molecular features were predicted based on *in-silico* mass fragmentation network analysis (SIRIUS) (*see* Methods; Supplementary Information Table 1). Principal component analysis (PCA) patterns were significantly different (PERMANOVA; F-value = 7.58; R^2^ = 0.62; adj. *p*-value=0.001) and showed that early stress (32°C) and recovery (27°C) metabolomes clustered together away from severe stress (34°C), suggesting that despite being separated by four months (June to October) the holobiont metabolic state during early stress and recovery were similar (Fig. 2a; Extended Data Fig. 1). The metabolomes of the diseased samples (L, AL) displayed inter-colony variations and showed a significant separation from all other samples, suggesting a fundamentally different chemical composition compared to healthy tissue (Fig. 2a). These findings confirm that acroporid corals in the Arabian Gulf exhibit a notable tolerance to temperatures up to 32°C, lethal to corals elsewhere, while exposure to 34°C poses a critical challenge to their overall fitness, which aligns with previous reports^28^.

**Figure 2.**
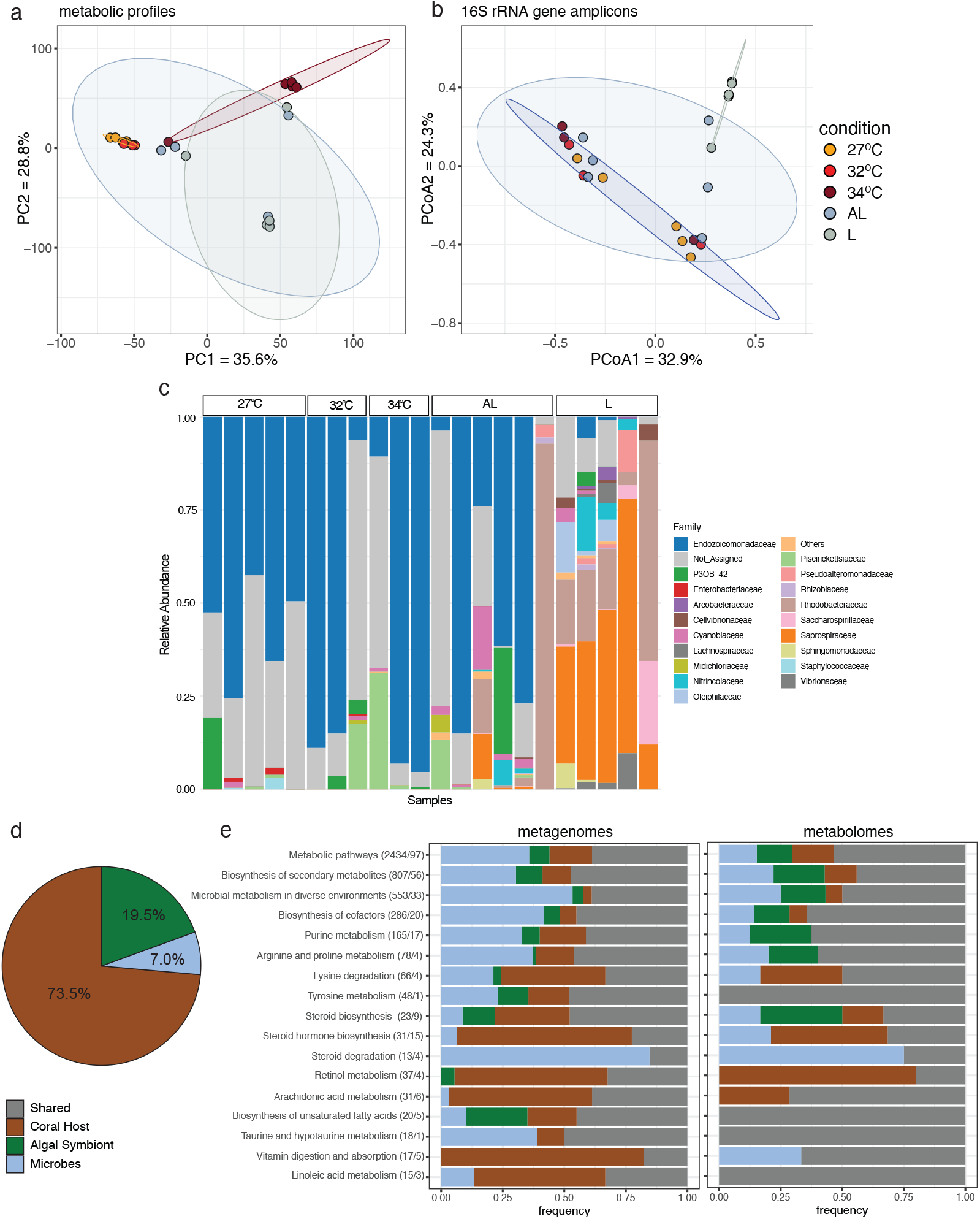
Integrated microbiome/metabolome analysis of the coral holobiont. **a**, Principal component analysis (PCA) of the metabolomics data (n=5878 putative molecular features). Samples collected at 27°C and 32°C cluster together and are significantly different from samples collected at 34°C. **b**, Principal coordinate analysis (PCoA) of microbial communities using Bray-Curtis distance shows significant differences between the diseased samples (i.e., AL and L) and all other samples. **c**, ASV-based bacterial community structure across temperatures and disease samples at the family level. Each bar plot represents the bacterial abundance of a coral colony. The top 20 bacterial families are shown. **d**, Distribution of metagenomic reads from coral holobiont metagenomes across coral, *Cladocopium spp*. (algal symbiont) and putative microbial members. **e**, Distribution of host-algal symbiont-bacterial genes (left) and metabolites (right) across the top-most abundant KEGG metabolic pathways. Numbers in parentheses refer to the number of genes and metabolites shown in each pathway, respectively.

The bacterial microbiome composition, based on 16S rRNA gene amplicons, revealed that disease samples (L, AL) differed significantly from all other samples (PERMANOVA; F-value = 1.98; R^2^ = 0.0318; *adj*. p-value = 0.01), suggesting the bacterial microbiome is resilient and only undergoes a significant shift during disease progression (Fig. 2b). Thirty-two bacterial ASVs belonging to the family Endozoicomonadaceae dominated the microbiome, with an average relative abundance of 60% in healthy samples. In addition, Endozoicomonadaceae persisted under elevated (32°C) and even extreme temperatures (34°C), which was particularly remarkable as they usually disappear during coral exposure to thermal stress^29^ (Fig. 2c; Extended Data Fig. 2a). At lesion sites, the relative abundance of Endozoicomonadaceae was only 1.3%, while Rhodobacteraceae and Saprospiraceae contributed 23% and 39% of the microbiome, respectively, indicating the proliferation of potentially opportunistic microbes at the expense of Endozoicomonadaceae during disease onset (Fig. 2c; Extended Data Fig. 2a)^30–32^.

### Integrated metagenome/metabolome analysis implicates the microbiome in steroid hormone metabolism

The metagenomes of coral holobionts produced a total of 589 gigabases (Gb), averaging 42 Gb per sample (Supplementary Information Table 3; Extended Data Fig. 3). We leveraged publicly available genomes for *Acropora* and *Cladocopium* to partition the holobiont metagenome before assembling coral and algal contigs (see Methods). A total of 257 million reads (∼7% of the holobiont reads) were classified as microbial (i.e., non-coral, non-algal reads; Fig. 2d). These microbial reads were used for downstream read-based and assembly-based metagenomic analyses (Extended Data Fig. 2b and 3; Supplementary Information Table 3). Taxonomic profiling of putative metagenomic microbial reads revealed a diverse coral microbiome primarily dominated by bacteria, comprising 95% of the microbial community (Extended Data Fig. 2b). To explore the potential roles of the microbiome, we assigned all genes and putatively annotated metabolites in our metagenomic and metabolomic datasets to members of the coral holobiont. To do this, metagenomic reads assigned to host, algal symbiont or bacteria were assembled and functionally annotated using the KEGG database to a total of 11,333 KEGG Orthology terms^33^ (KOs). While a third of the predicted KOs (3,065) were shared between different members of the holobiont, most were uniquely assigned to the coral host (4,619), followed by microbes (3,029) and algal symbionts (620) (Fig. 2e; Supplementary Information Table 4). Using these KO taxonomic assignments, we mapped the putatively annotated metabolites associated with KEGG compounds (284) (Supplementary Information Table 2). This analysis indicated 148 of these putative metabolites were shared between different members of the holobiont, while 43 were assigned uniquely to the host, 42 to the algal symbiont and 51 to other microbes (Fig. 2e, Extended Data Table 2). Examining the KEGG pathways across metagenomes and metabolomes indicated that the host dominates the biosynthesis of steroid hormones, retinol metabolism, and vitamin digestion and absorption, while microbes dominate biosynthesis of cofactors, biosynthesis of secondary metabolites, and steroid degradation (Fig. 2e).

To assign functions to specific bacterial taxa, we assembled genomes from our metagenomes (MAGs). Coral samples devoid of mucus and seawater yielded 11 medium to high-quality MAGs (completeness >75% and contamination <10%). Core microbiome members were defined as MAGs consistently present at relatively high (40-60%) abundance in healthy tissues (at all temperatures), but showing reduced abundance in dysbiotic samples (AL and L) (Fig. 3a). Out of the 11 MAGs, two MAGs stood out as important members of the coral holobiont: MAG01 assigned to the phylum Myxococcota and MAG10 assigned to the genus *Endozoicomonas* (Extended Data Fig. 4; Supplementary Information Table 5). *Endozoicomonas* (MAG10; completeness: 91%, contamination: 1.4%), exhibited a dynamic abundance profile as its relative abundance peaked at 32°C during early stress (65.4%), followed by a dramatic decrease at 34°C (35.8%) and then bounced back with a modest abundance during recovery at 27°C (43%) (Fig. 3a). The relative abundance of MAG10 dropped drastically in L samples (0.8%), coinciding with dysbiosis and consistent with previous reports^30–32^. Given its dynamic abundance pattern and strong association with healthy coral states, we next explored the functional potential of MAG10 to better understand its role within the coral holobiont.

**Figure 3.**
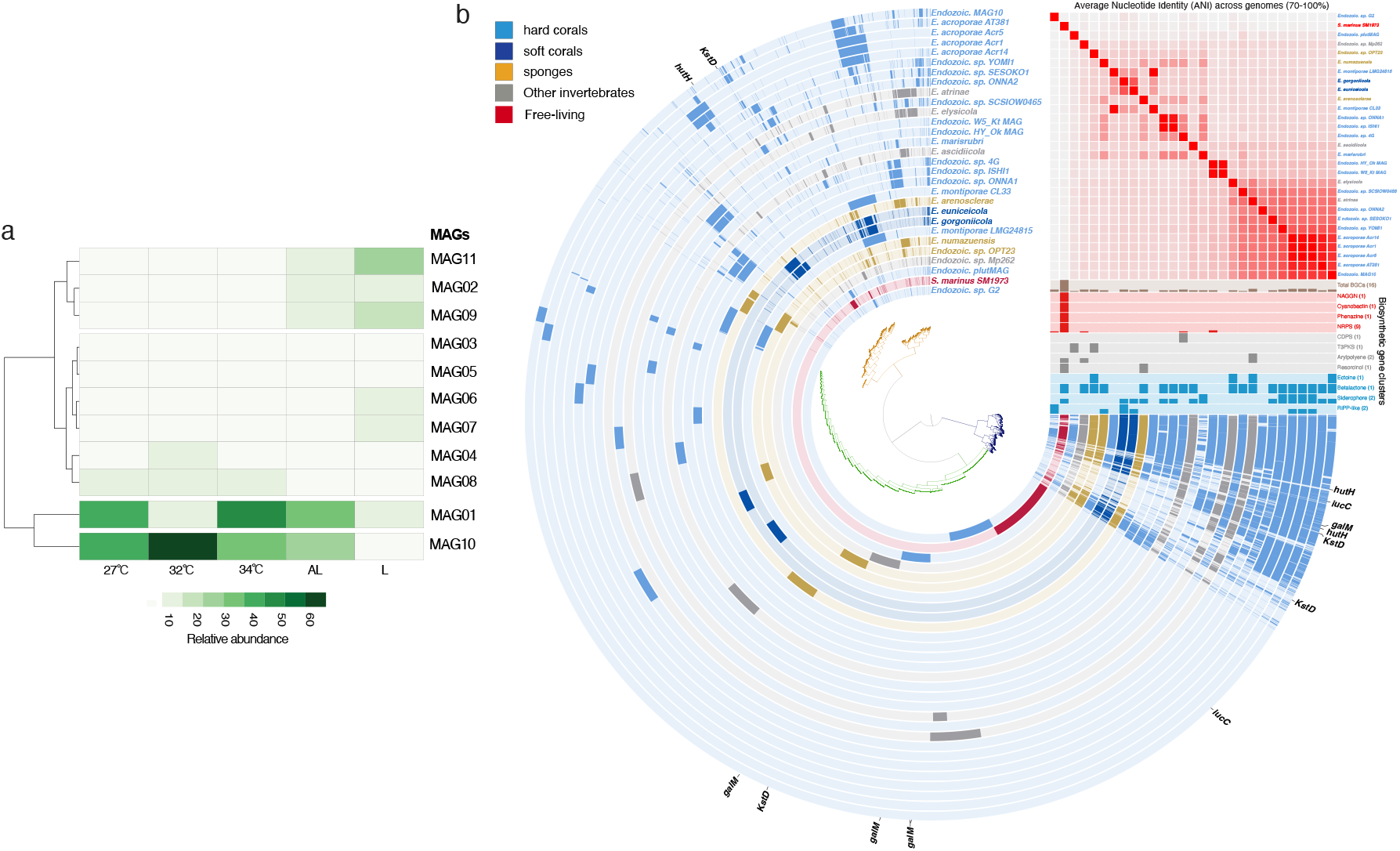
Abundance of *Endozoicomonas* MAG10 and the putative functional importance of *Endozoicomonas*. **a**, Relative abundance of tissue-associated MAGs inferred by read recruitment from metagenomes across all samples. Taxonomic classifications of MAGs are shown in SI Table 5. **b**, *Endozoicomonas* pangenome and biosynthetic gene clusters (BGCs) of 29 genomes and MAGs, including MAG10 and *E. acroporae* AT381. *Spartinivicinus marina* SM1973 was included to represent a free-living member of the Oceanospirillales as a close free-living isolate to *Endozoicomonas*. Pangenomic analyses based on the occurrence of gene clusters were implemented in Anvi’o^34^. The central dendrogram represents hierarchical clustering based on the presence (colored) or absence (opaque) of 34,846 gene clusters. Branches were colored based on core, accessory or singleton/doubleton genes. Each track in the pangenome represents a single genome colored according to its isolation source. Genomes were clustered based on their pairwise average nucleotide identity (ANI) values shown in the heatmap, which reveals high similarity between MAG10 and *E. acroporae* genomes. Distribution of BGCs across genomes is shown below the heatmap. Key genes related to aerobactin biosynthesis (*iucC*) and degradation of steroid hormones (*kstD*), galactose (*galM*), and histidine (*hutH*) are labeled on the periphery of the pangenome.

### Pangenome analysis reveals conserved functions across *Endozoicomonas* genomes

Since culturing efforts of *Endozoicomonas* from *Acropora* in the Arabian Gulf were not successful, we isolated *Endozoicomonas acroporae* AT381 from *A. kenti* from the Great Barrier Reef (GBR), referred to hereafter as AT381 (Fig. 1b), and obtained a nearly complete genome (size = 6.02 Mbp; completeness = 99.1%) predicted to harbor 5001 coding sequences (CDS). AT381 and MAG10 are closely related as they share an average nucleotide identity (ANI) of 94.3%^34^. Phylogenomic analysis between MAG10, AT381 and 27 publicly available *Endozoicomonas* genomes and MAGs from diverse invertebrate hosts further confirmed the close relationship between MAG10, AT381 and other *E. acroporae* genomes (Extended Data Fig. 5). Functional conservation and specialization across all *Endozoicomonas* genomes were examined by constructing a pangenome of their 34,846 gene clusters (Fig. 3b, Supplementary Information Table 6)^35^. Because of the inability to culture a MAG10 relative from the Arabian Gulf, we used pangenomic analysis to confirm functional conservations within the *Endozoicomonas* genus and further functionally validated some of these conserved capabilities using the closely related *E. acroporae* AT381 from the GBR and two other *Endozoicomonas* isolates.

We hypothesized that *Endozoicomonas* genomes symbiotic with corals possess specialized, but shared functions enabling interactions with their coral hosts, compared to genomes isolated from other hosts. As reported for other genomes of coral-associated *Endozoicomonas* spp., MAG10 harbors genes related to symbiosis establishment, e.g., flagellar assembly, type IV pili^18^, ankyrin and tetratricopeptide repeats^21,22^, several secretion systems involved in host infection (T2SS, T3SS, T6SS)^18,21–23^, complete biosynthesis pathways for the vitamins riboflavin (B_2_), biotin (B_7_) and pyridoxal-phosphate (B_6_), and nine essential amino acids and their transporters (Extended Data Fig. 4b and c, Supplementary Information Table 7)^11^. MAG10 was also significantly enriched within the holobiont for antioxidant production^22^, heme biosynthesis^25^, glycoside hydrolases that cleave the most abundant marine polymers, starch and chitin^36^, and diverse ABC transporters (Supplementary Information Tables 7 and 8). In addition to these previously known functions, MAG10 is capable of **(a)** catabolizing histidine to glutamate, an essential intermediate for ammonium assimilation and a common feature of most *Endozoicomonas* genomes, **(b)** degrading galactose (Leloir pathway), a Symbiodiniaceae photosynthate^37^, which is a common feature in most *Endozoicomonas* genomes, **(c)** biosynthesizing the nucleoside antibiotic showdomycin, **(d)** biosynthesizing and exporting the polyamine spermidine, a reactive oxygen species scavenger, and export proteins for its precursor putrescine, **(e)** and biosynthesizing ectoine, a bacterial osmoprotectant (Fig. 3b, Extended Data Fig. 4c, Extended Data Tables 3 and 4, Supplementary Information Table 7). Cumulatively, these functions may enhance symbiosis and increase the fitness of the coral host and/or Symbiodiniaceae, especially during thermal stress.

### *Endozoicomonas* acquires iron *in vitro* using the siderophore aerobactin

Siderophores are small molecules secreted by some bacteria and fungi to acquire iron in iron-limited environments^38^ and can play important roles in iron cycling and bioavailability in the pelagic ocean^39,40^. The genome of *Klebsiella pneumoniae*, a model bacterium for studying the biosynthesis of the siderophore aerobactin, possesses the canonical, complete gene cluster/operon containing aerobactin synthases (*iucA/iucC*) and the TonB-dependent uptake receptor (*iutA*) used to produce apo-aerobactin and take up Fe-bound aerobactin, respectively^41^. In contrast, MAG10, all other *E. acroporae* and 8 additional coral- and invertebrate-associated *Endozoicomonas* genomes possessed only the biosynthesis genes within the gene cluster, while having either a distal *iutA* that is outside of the gene cluster in the case of AT381 or a homolog of *iutA* in a completely separate region of the genome for all other *Endozoicomonas* that possess the gene cluster (Fig. 4a, Extended Data Table 4). Aerobactin requires a functional *iutA* for Fe acquisition to function^42^ and a distal *iutA* suggests a potential decoupling from biosynthesis. To confirm whether *Endozoicomonas* can use aerobactin to acquire iron, we grew multiple strains *in vitro* under Fe-limiting conditions and tested their siderophore production using the chrome azurol S assay (CAS)^43^. The tested strains include two isolates from *Acropora*: *E. acroporae* AT381 and *E. acroporae* Acr14, and *E. montiporae* CL33 (isolated from *Montipora*), a negative control that lacks aerobactin biosynthesis genes (Extended Data Table 4). The CAS assay showed that *E. acroporae* AT381 and Acr14 were able to grow and utilize Fe from CAS (Fig. 4b) while *E. montiporae* was not able to grow, consistent with its inability to produce aerobactin. These findings suggest that select *Endozoicomonas* strains and species (12 out of 27 in total) can utilize siderophores for Fe acquisition *in vitro* and likely *ex hospite*. We were not able to detect any m/z signatures that match aerobactin, its Fe-complex or photoproducts^44^ in the holobiont metabolomes. Indeed, siderophore production *in hospite* is mainly associated with pathogenic bacteria^45^, and siderophore production may take place when *Endozoicomonas* are in seawater *ex hospite*.

**Figure 4.**
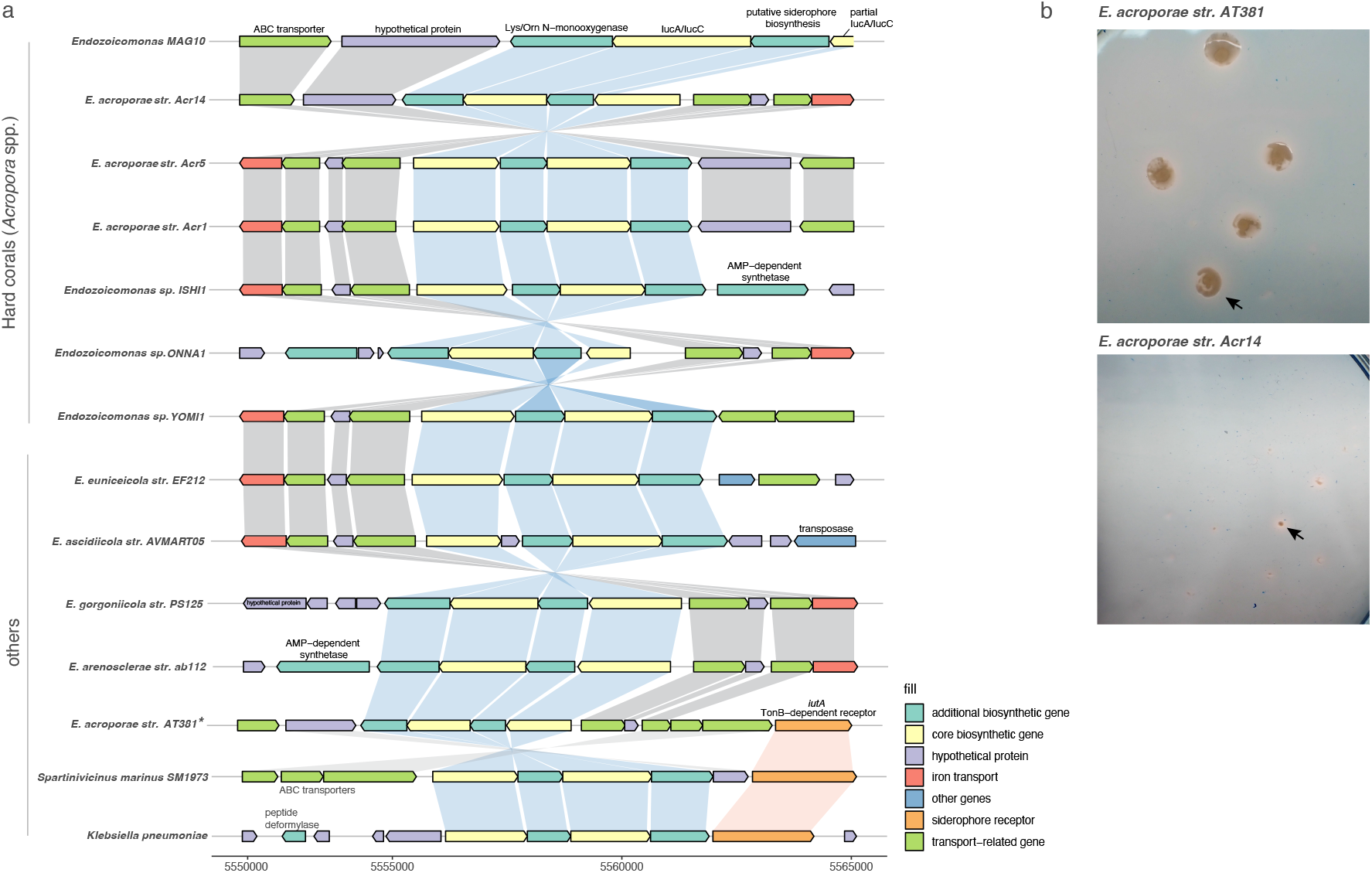
Utilization potential of the siderophore aerobactin for iron acquisition in *Endozoicomonas*. **a**, Genomic neighborhood organization of the aerobactin BGC in genomes of MAG10, *Endozoicomonas* spp. symbiotic with *Acropora* corals, soft corals, their closest free-living isolate, *S. marinus* SM1973, and the aerobactin-producer *Klebsiella pneumoniae*. Shading across genes represents gene synteny of aerobactin biosynthesis (blue), aerobactin uptake (orange), or other genes (grey). Aerobactin is biosynthesized by 4 genes, one of which is duplicated (*iucA/iucC*). Although the aerobactin biosynthesis operon is partially truncated in the MAG10 scaffold, we confirmed that the only partially truncated gene is the duplicated *iucA/iucC*. The TonB-dependent receptor refers to the aerobactin transporter gene, *iutA*, which is proximal to the BGC of SM1973 and K. *pneumoniae*. Among all coral-associated *Endozoicomonas*, only *E. acroporae* AT381 has *iutA* ∼11kb distal to the BGC; all other *Endozoicomonas* genomes display a distal ortholog/paralog of *iutA* elsewhere in the genome (>11kb away from the biosynthesis BGC) **b**, CAS assay of cultures of two *E. acroporae* strains (AT381 and Acr14), encoding the aerobactin cluster, grown in iron-limited media on agar. Siderophore activity was then assessed through the formation of halos (black arrow) around bacterial colonies. * indicates isolation from hard coras from the genus *Acropora*.

The fact that all examined *Endozoicomonas* genomes capable of biosynthesizing aerobactin possess distal, not proximal, *iutA* homologs indicates a potential decoupling of expression between biosynthesis and transport, which can make Fe acquisition less efficient and more energetically costly for the bacteria. This pattern also suggests a gradual loss of function of the transporter that would eventually abolish aerobactin biosynthesis over evolutionary timescale. Indeed, a loss of function was concluded from the lack of *iutA* in the genomes of *Bacillus licheniformis* and *B. halodurans* despite the presence of the aerobactin biosynthesis operon^46^.

To assess the evolutionary pressures acting on the *iutA* gene, we calculated rates of non-synonymous and synonymous substitutions (dN/dS) ratios across multiple *E. acroporae* strains compared to the housekeeping gene DNA Polymerase III subunit delta as a conserved core gene against *E. acroporae* AT381 as a reference (Supplementary Information Table 9). Several *Endozoicomonas* isolates from *Acropora* displayed signatures of relaxed purifying selection (significantly elevated dN/dS) in *iutA*, suggesting ecological redundancy or diminished reliance on siderophore-mediated iron acquisition in certain niches. In contrast, *iutA* in *E. acroporae* Acr14 exhibited low dN/dS value (0.1), consistent with strong purifying selection and functional conservation. These patterns suggest that while some *Endozoicomonas* strains may reduce dependence on siderophore systems, others retain this capability, potentially as a survival strategy during transitions to a free-living lifestyle *ex hospite*. MAG10 was not included in this analysis because MAGs are typically a mosaic of sequences from multiple closely related strains, rather than a single, clonal genome. This chimerism, along with assembly artifacts, can distort substitution rate estimates and obscure true evolutionary signals^47^, a concern that is particularly relevant here given the high strain-level diversity we observed, with 32 distinct ASVs from Endozoicomonadaceae identified by 16S rRNA gene amplicon sequencing. Thus, we propose that coral-associated *Endozoicomonas* do not utilize siderophore-mediated iron uptake *in hospite*, a function that could be used in the free-living stage, and that over evolutionary time aerobactin-dependent Fe acquisition will be lost. Coral-associated *Endozoicomonas* can instead utilize other pathways to acquire iron, such as heme transporters (Supplementary Information Table 7).

### Steroid metabolism is a prominent function of *Endozoicomonas* among the coral microbiome

MAG10 was the only holobiont member that possessed the complete pathway to utilize carbon from the degradation of the steroid hormone testosterone via 3-oxosteroid 1-dehydrogenase (*kstD*) gene (Fig. 5a,b; Supplementary Information Table 4, 7; Extended Data Fig. 3). This function was shared with most *Endozoicomonas* genomes (Fig. 3b; Extended Data Table 4), consistent with their broader capacity to break down this important steroid molecule^48^. The relative abundance of most steroids gradually and significantly decreased as temperature increased, with abundance at 34°C, AL, and L near background levels (Fig. 5a, Extended Data Fig. 6, Supplementary Information Table 9), consistent with previous reports^49,50^ showing suppression of steroid hormone production under stress and during infection. Surprisingly, a notable exception to this pattern were select key terminal steroids and steroid degradation products. Specifically, the putative eukaryotic hormones, progesterone, testosterone and estradiol-17β, in addition to *Endozoicomonas*-derived androsta-1,4-diene-3,17-dione and HIP showed the highest relative abundance at 34°C, with most other samples at near background levels (Fig. 5a, Supplementary Information Table 10). Since the sampling timepoints fell outside the spawning season in the Arabian Gulf (March to May)^51^, and most precursors of progesterone and testosterone showed low relative abundance near background levels at 34°C, the elevated levels of these putative steroid hormones during severe stress at 34°C were unexpected. Since these precursors are required for the synthesis of testosterone and progesterone, their absence suggests they are either depleted by the host or a source other than the host, is providing testosterone and progesterone. Notably, MAG10 was the only microbiome member that possessed a 17-β-hydroxysteroid dehydrogenase (K05296; E1.1.1.51) that synthesizes testosterone from androstenedione, the reverse reaction of KstD (Supplementary Information Table 7)^52^. Therefore, we hypothesized that *Endozoicomonas* may provide corals with some of these steroids during severe stress.

**Figure 5.**
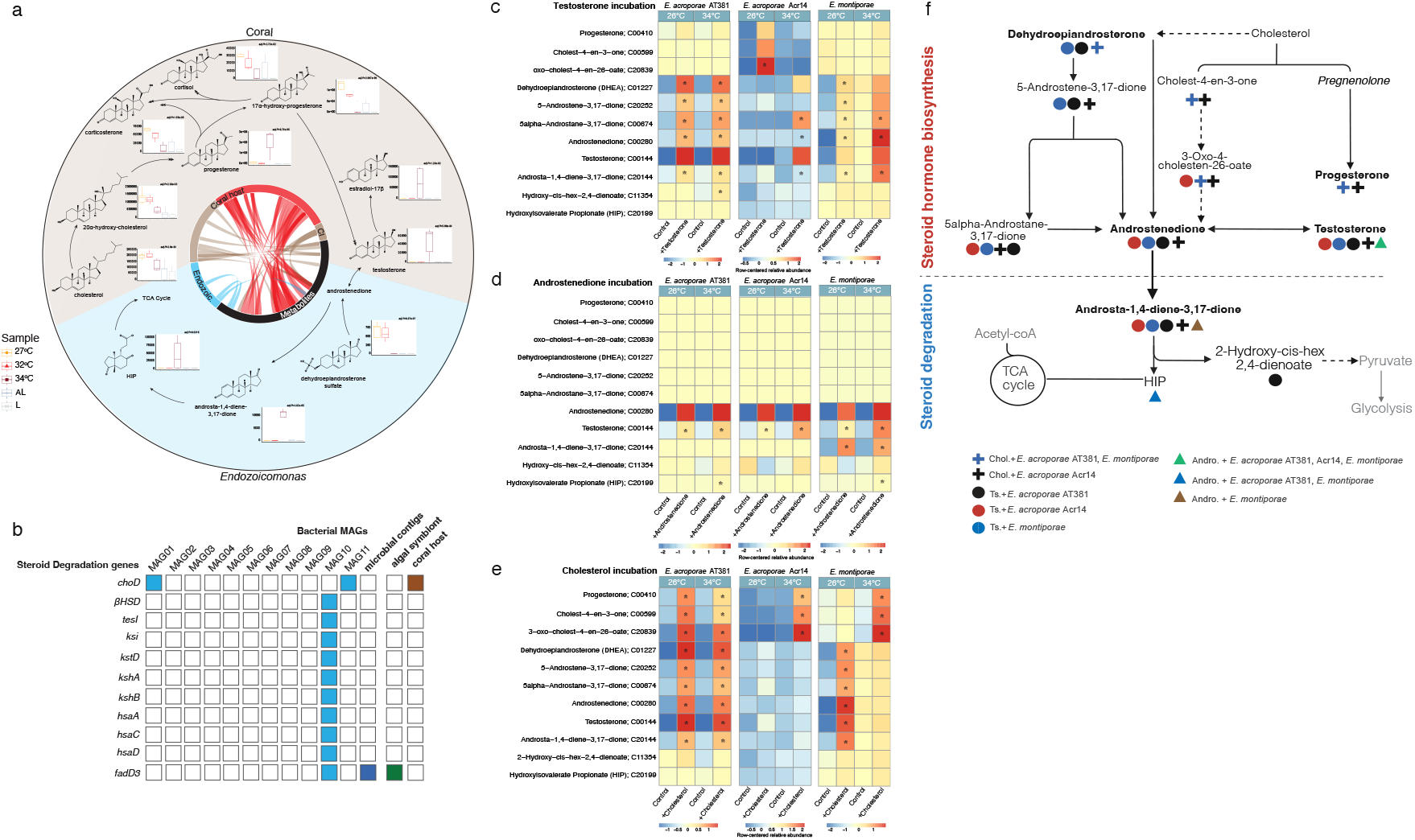
Putative steroid hormone biosynthesis/degradation in the holobiont across the temperature gradient and an abbreviated steroid hormone production pathway by *Endozoicomonas*. **a**, Steroid biosynthesis and degradation pathways based on coral and Endozoicomonas MAG10 contigs. Arrows indicate genes, and arrowheads correspond to the number of genes carrying a transformation. The relative abundance of putative steroid hormones is presented along with adjusted p-values, calculated using the Benjamini-Hochberg method, based on ANOVA results that assess the impact of temperature on changes in their abundance. Middle: Circos plot depicting all detected steroid-related metabolites and their connection to genes of *Endozoicomonas* (Endozoic.), the coral host, or *Cladocopium* spp. (Cl). The plot shows that steroid degradation is restricted to *Endozoicomonas*, steroid biosynthesis is mostly dominated by *Cladocopium* and steroid hormone biosynthesis is mostly dominated by the coral. Colors indicate different pathways (brown=steroid biosynthesis, red=steroid hormone biosynthesis, blue=steroid degradation, black=metabolites). **b**, Presence/absence of steroid degradation genes in the holobiont. Note that microbial contigs excluded contigs used to construct the bacterial MAGs. cholesterol oxidase; choD, beta-hydroxysteroid dehydrogenase; βHSD, 3-oxo-5alpha-steroid 4-dehydrogenase; tesI, steroid Delta-isomerase; ksi, 3-oxosteroid 1-dehydrogenase; kstD, 3-ketosteroid 9alpha-monooxygenase subunit A; kshA, 3-ketosteroid 9alpha-monooxygenase subunit B; kshB, 3-hydroxy-9,10-secoandrosta-(10)1,3,5-triene-9,17-dione monooxygenase; hsaA, 3,4-dihydroxy-9,10-secoandrosta-(10)1,3,5-triene-9,17-dione 4,5-dioxygenase; hsaC, 4,5:9,10-diseco-3-hydroxy-5,9,17-trioxoandrosta-2,(10)1-diene-4-oate hydrolase; hsaD, HIP-CoA ligase; fadD3. **c, d and e**, Steroid hormone production by three *Endozoicomonas* strains. Two *E. acroporae* (AT381 and Acr 14) and one *E. montiporae* strains were incubated with or without testosterone (**c**), Androstenedione (**d**), or cholesterol (**e**), at 26°C and 34°C. Heatmaps show the normalized mean relative abundance of each metabolite in each condition, with * indicating statistical significance of their treatment relative to controls (T-test, adjusted p-value <0.05). Statistical significance was not calculated for testosterone/androstenedione in **c, d** because both were added to the media. All detected metabolites were annotated based onaccurate mass within 5 ppm and high-resolution MS/MS fragmentation of standards. Additionally, a testosterone standard was used to further confirm its production using the aforementioned criteria and retention time. **c**, Novel pathway for the production of progesterone and testosterone inferred from metabolomics analysis. Dotted lines indicate the absence of genes involved in these specific metabolic reactions in the *Endozoicomonas* genomes.

MAG10 and other *Endozoicomonas* genomes (e.g., *E. montiporae*) encode genes associated with cholesterol binding, such as those containing the steroidogenic acute regulatory protein-related lipid transfer (START) and sterol carrier protein-2 (SCP2) domains, as well as cholesterol/phospholipid transport, including MlaD, MlaE, and ABC transporters (Supplementary Information Table 7). Therefore, we investigated whether *Endozoicomonas* can catabolise testosterone, convert androstenedione to testosterone and if it synthesizes the terminal steroids progesterone, testosterone, or estradiol-17β from cholesterol, given that cholesterol is the precursor of steroid hormones and can be one of the abundant steroid molecules in the coral holobiont^48,53^. Cultures of three *Endozoicomonas* strains, *E. acroporae* AT381, *E. acroporae* Acr14 and *E. montiporae* were grown at 26°C and 34°C in minimal media with or without cholesterol, testosterone or androstenedione (Fig. 1b). After 24 hours, all three strains catabolized testosterone into androstenedione and downstream metabolites leading to the TCA cycle, with *E. acroporae* Acr14 only carrying out this transformation at 34°C, while *E. acroporae* AT381 and *E. montiporae* doing so at both temperatures (Fig. 5c, Supplementary Information Table 11), suggesting thermal regulation of these transformations in *E. acroporae* Acr14. All three strains were also able to convert androstenedione to testosterone at both temperatures likely via 17-β-hydroxysteroid dehydrogenase (Fig. 5d, Supplementary Information Table 11). Surprisingly, all three strains were capable of transforming cholesterol into terminal steroids. *E. acroporae* AT381 converted cholesterol into the terminal products progesterone, testosterone, androstenedione and other intermediates at both temperatures (Fig. 5e, Extended Data Fig. 6c). Conversely, *E. acroporae* Acr14 converted cholesterol to cholesterol derivatives and progesterone only at 34^°^C with no transformations observed at 26^°^C. Finally, *E. montiporae* converted cholesterol to androstenedione, testosterone and related metabolites only at 26^°^C and switched to converting cholesterol to cholesterol derivatives and progesterone at 34^°^C, indicating a strong temperature influence on the regulation of these transformations (Fig. 5e). These findings suggest that depending on the species/strain of *Endozoicomonas*, cholesterol conversion towards testosterone or progesterone is controlled by temperature, which may explain the high relative abundance of progesterone and testosterone in the holobiont at 34^°^C (Fig. 5 and Extended Data Fig. 6a,b).

Using high-resolution mass spectrometry and fragmentation information, we propose a metabolic pathway that bifurcates cholesterol into **(a)** progesterone via pregnenolone and **(b)** testosterone via oxo-cholestenoate and androstenedione (Fig. 5f). Although the *Endozoicomonas* pangenome and auto-annotation highlight important enzymes for steroid degradation (e.g. *kstD*), further mining of the *E. montiporae* genome yielded steroid delta isomerases of the CyP450 family, which putatively convert pregnenolone into progesterone (Fig. 5f; Supplementary Information Table 7)^48^. MAG10 also encodes a member of the 3-beta hydroxysteroid dehydrogenase/isomerase protein family [PF01073.22] (Supplementary Information Table 7). Orthologs of this protein are present in all three *Endozoicomonas* strains we used for steroid incubations. This enzyme family includes 3β-hydroxysteroid dehydrogenase/Δ^5^-Δ^4^ isomerases (3β-HSDs), which plays a central role in steroid biosynthesis by catalyzing the oxidation and isomerization of steroid precursors into active steroid hormones. Although mining *Endozoicomonas* genomes did not reveal any specific exporters for testosterone or progesterone, both hormones are lipophilic and can diffuse freely through the cytoplasmic membrane^54–56^. Collectively, these findings indicate that *Endozoicomonas* possess the genetic repertoire not only for steroid degradation, but for their conversion too, positioning them as active contributors to steroid metabolism in the coral holobiont and suggesting that these steroids play an integral role in the coral response to thermal stress. Because testosterone is an immunosuppressive substrate under stress and progesterone is an immunomodulatory and anti-inflammatory substrate in various animal models, we hypothesized that the production of these hormones by *Endozoicomonas* likely modulates how the coral immune system responds to thermal stress.

### Steroid hormones prime the coral immune response during thermal stress

To determine whether corals can take up exogenous steroid hormones and their effective concentrations, we incubated *Acropora* nubbins with three different concentrations of ^13^C-labeled testosterone (10 nM, 100 nM, 1 *μ*M) for 48 hours and examined its relative abundance after 24 and 48 hours (Fig. 6a). While corals incubated with 1*μ*M accumulated the most testosterone after 24 hours, the steroid relative abundance was highly reduced (3-fold) at 48 hours, suggesting saturation. Similar results were obtained with incubations with 100 nM testosterone. At 10 nM, *Acropora* accumulated testosterone progressively at 24 and 48 hours, resulting in an 11-fold increase indicative of active uptake and limited metabolic removal, consistent with a potential biological role; therefore, we used this concentration for subsequent experiments for both testosterone and progesterone. To test the effect of each molecule on the immune response of *Acropora*, coral nubbins were incubated with each molecule for 3 hours at an ambient temperature of 28°C (T1), followed by an increase in temperature to 33°C (T2), simulating thermal stress and at which point a second dose of progesterone or testosterone were added. To map the coral immune system response to these steroids during thermal stress, samples were collected at T1, T2 and at 24 hours post T1 (T3) (Fig. 6b) for RNA sequencing. Coral fragments were monitored by measuring algal symbiont’s photosynthetic efficiency (*Fv/Fm)* to ensure corals were stable and exposure to both steroid hormones did not lead to physiological toxicity (Fig. 6c).

**Figure 6.**
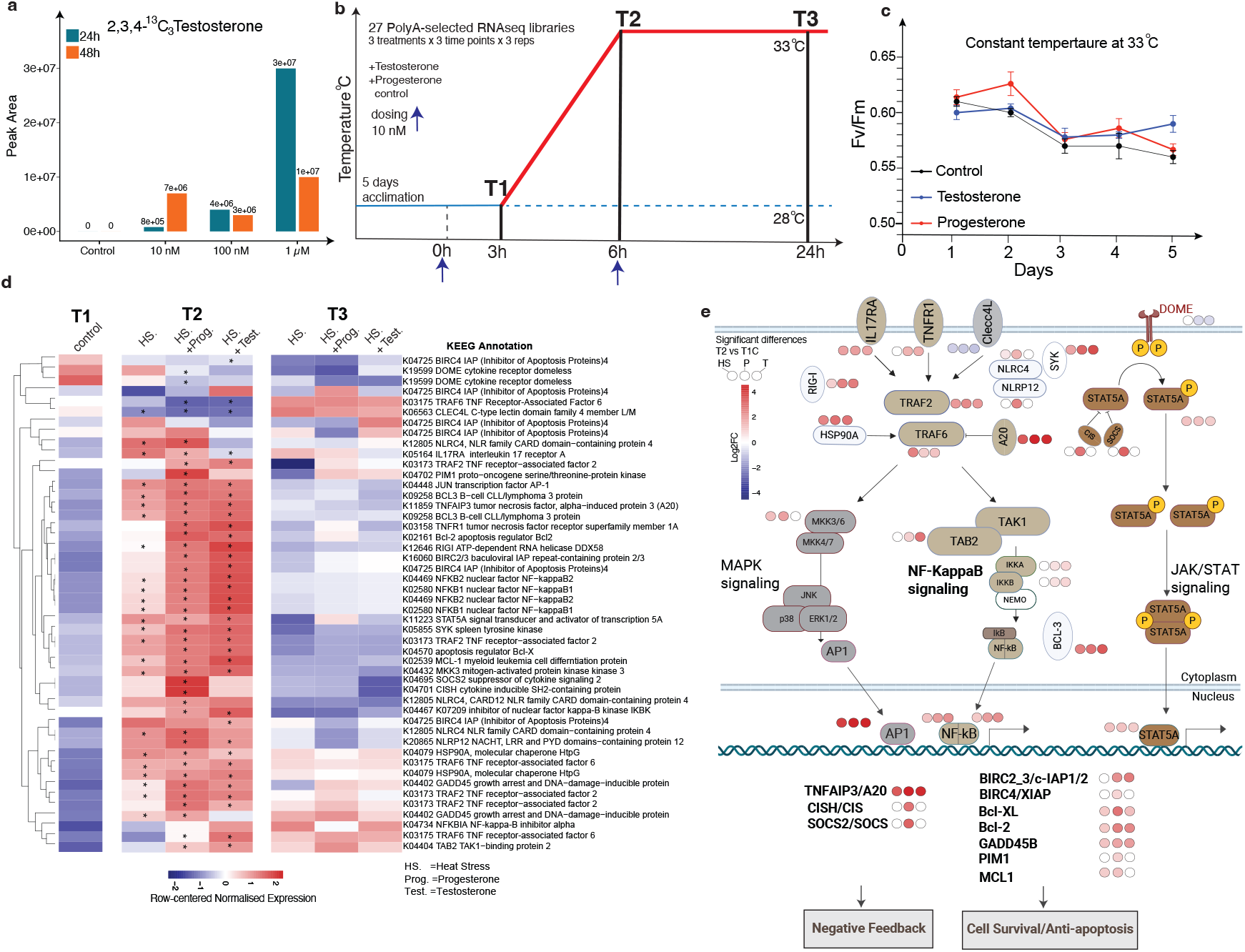
Activation of the coral innate immune response by progesterone and testosterone during heat stress. **a**, Uptake and accumulation of ^13^C-labeled testosterone in coral tissues over a 48-h exposure period. Peak area was normalised to dry extract weight. **b**, Experimental design. *Acropora* colonies were exposed to acute thermal stress with or without the addition of progesterone or testosterone. Hormones were administered at 10 nM at the start of the experiment at 28°C. Three hours later, the first sampling point (T1) was collected for RNA-seq analysis, followed by a second hormone dose at 6 h. The temperature was ramped up from 28°C to 33°C over 3 h after T1 collection, and samples were collected at T2 along with untreated controls. Final sampling (T3) was conducted after 24 h of exposure at 33°C. **c**, Maximum quantum yield of photosystem II (F_v_/F_m_) was measured daily as an indicator of photophysiological stress and a proxy for the coral holobiont health. **d**, Heatmap showing normalized expression levels of selected immunity-related genes differentially regulated across treatments and time points. Significantly differentially expressed genes at 6 hours compared to controls (FDR < 0.05) were highlighted by asterisks. All differential expressions were based on biological triplicates. **e**, Pathway diagram based on KEGG annotations highlighting genes associated with innate immune pathways, including immune receptors, intracellular signaling cascades, and downstream effector responses. Tumor necrosis factor receptor 1 (TNFR1), TNF receptor–associated factor 2 (TRAF2), TNF receptor–associated factor 6 (TRAF6), TAK1-binding protein 2 (TAB2), Mitogen-activated protein kinase kinase 3 (MKK3), Mitogen-activated protein kinase kinase 4/7 (MKK4/7), Nuclear factor kappa-B subunit 1/2 (NFKB1/2), NF-κB inhibitor alpha (IκBα / NFKBIA), Inhibitor of nuclear factor κB kinase subunit alpha/beta (IKBKA/B), NF-κB essential modulator (NEMO / IKBKG), Activator protein 1 (AP-1), C-type lectin domain family 4 member L (CLEC4L), Retinoic acid–inducible gene I (RIG-I), Interleukin-17 receptor A (IL17RA), Signal transducer and activator of transcription 5A (STAT5A), Cytokine receptor domeless (DOME), Spleen tyrosine kinase (SYK), B-cell lymphoma 2 (*BCL2*), B-cell lymphoma extra-large (*BCL-XL*), baculoviral IAP repeat-containing proteins 2 and 3 (*BIRC2/3*), tumor necrosis factor alpha–induced protein 3 (*TNFAIP3*), cytokine-inducible SH2-containing protein (*CISH*), suppressor of cytokine signaling 2 (*SOCS2*), heat shock protein 90 alpha (*HSP90A*).

To examine differential expression of immune-related genes upon coral exposure to testosterone or progesterone during thermal stress relative to untreated controls (Supplementary Information Table 12, 13, 14), we compiled a list of all known innate immunity-related genes from our assembled transcriptome (Fig. 6d). While many immune-related genes were activated due to heat stress, exposure to testosterone or progesterone further increased their transcription (Fig. 6d,e). For example during T2, both testosterone and progesterone treatments caused on average an increase in the fold-change expression of master regulators of innate immunity, including tumor necrosis factor receptor (TNFR; +129%), nuclear factor kappa B (NF-κB; +80%), and activator protein 1 (AP-1; +10%) relative to controls, indicating increased immune signal transduction and immune priming. Concomitantly, upstream immune-sensing and adaptor pathways (e.g., RIG1 and *TRAF2*; +131%, +1.9% respectively), as well as downstream genes linked to immune regulation and anti-apoptosis (e.g., *TNFAIP3, BIRC2/3, BCL2*; +18.5%, +64%, +104% respectively) were activated. This suggests that both testosterone and progesterone prime the coral innate immune system during thermal stress by enhancing immune signaling and activating both upstream sensing and downstream protective genes. Expression of pro-inflammatory signaling components decreased (*TRAF6*; −32% and the JAK/STAT receptor *DOME*; −84%), indicating immune-dampening that limits immune overreaction. Despite these shared effects in the presence of testosterone and progesterone, progesterone uniquely enhanced transcriptional responses associated with cell survival and anti-apoptotic signaling (*MCL1, BIRC4, PIM1*; +25%, +161%, +245% respectively) and preferentially promoted negative feedback regulation of cytokine-activated JAK/STAT pathway (*CISH/SOCS2*; +190%), consistent with suppression of prolonged or excessive immune activation (Fig. 6d,e). Coincidentally, progesterone was preferentially produced by some *Endozoicomonas* from cholesterol only during temperatures resembling thermal stress (Fig. 5e), suggesting *Endozoicomonas* modulates the coral immune system to suppress prolonged inflammatory responses to thermal stress.

## Discussion

Despite the central role of microbial associates to coral health and fitness, the functional mechanisms bacteria utilize to influence host physiology remain largely unresolved. Here, we provide the first evidence that ubiquitous *Endozoicomonas* possesses an unexpected capacity to synthesize terminal eukaryotic steroid hormones, a novel metabolic capability in the coral microbiome, and that these hormones modulate the host immune response during thermal stress, suggesting a critical functional role for this genus within the coral holobiont. Particularly, the ability of *Endozoicomonas* to synthesize testosterone and progesterone in a temperature-dependent manner confirms that their accumulation in the holobiont when seawater temperature is high may be partially or wholly driven by *Endozoicomonas* symbionts. While steroid degradation is widespread in many alpha- and gamma-proteobacteria^48,57,58^, the ability to synthesize terminal, host-active steroid hormones from cholesterol is rare and positions *Endozoicomonas* as a metabolic outlier among coral symbionts. Specific human gut microbiota, such as *Bacteroides acidifaciens* and *Ruminococcus gnavus*, synthesize steroid hormones and play a major role in steroid hormone homeostasis in mammals^59^. Moreover, *Mycobacterium smegmatis* was shown to synthesize testosterone and estrogens *in vitro* from steroid precursors^60^. While the mycobacterial conversion of cholesterol to testosterone has been observed under fermentative conditions, *Endozoicomonas* differs in its ability to carry these transformations under aerobic conditions, highlighting a distinct biochemical strategy that can influence major functions in their host.

While steroid hormones play critical roles in many physiological functions, such as growth, reproduction, and development in vertebrates^48^, their role and origin in invertebrates have been subject to recent debate^61^. While definitive CYP19A1 and HSD17B3 orthologs (the vertebrate enzymes that catalyze the final steps in estrogen and androgen synthesis, respectively) have not been identified in invertebrate genomes, several studies have demonstrated functional aromatase activity in corals (e.g., *Euphyllia ancora, Pocillopora damicornis*) using substrate conversion assays and inhibition by vertebrate aromatase inhibitors^62,63^ that suggest invertebrates are capable of synthesizing these molecules. We also identified several coral genes involved in steroid hormone biosynthesis (Supplementary Information Table 7), including 17β-hydroxysteroid dehydrogenase and several members of the cytochrome P450 enzyme family that can biosynthesize testosterone and progesterone, respectively, as reported in previous coral genomes^64^. This suggests that although the exact gene homologs that produce steroid hormones may differ, the enzyme function is present.

In corals, steroid hormones play important roles in gametogenesis and spawning and exogenous additions of steroid hormones influence the coral microbiome composition^61,65,66^. Our findings indicate that, in addition to these functions, steroid hormones modulate the immune response to thermal stress within hours of progesterone or testosterone addition. Specifically, hormone exposure under thermal stress preferentially engaged TRAF2-associated TNF/NFKappaB signaling and upregulated immune negative regulators, suggesting a survival-oriented, immune-modulating state rather than a canonical inflammation response. TRAF signaling can diverge toward either pro-survival or pro-inflammatory outcomes, with TRAF2 typically implicated in cell survival and anti-apoptotic signaling^67^ and TRAF6 commonly associated with inflammatory pathways^68^. Indeed, exposure to either progesterone or testosterone under thermal stress induced TRAF2 expression while suppressing TRAF6, suggesting a shift from inflammatory to anti-apoptotic response relative to the coral’s response under thermal stress alone (Fig. 6e). In contrast to the combined steroid hormone response, progesterone uniquely attenuated the JAK/STAT signaling pathway by downregulating DOME receptors and inducing the negative regulators of JAK/STAT, CIS and SOCS (Fig. 6e). These patterns may fine-tune the immune response by limiting immune overactivation while preserving basal immune competence^69,70^. By priming innate immunity while constraining excessive inflammation, progesterone may help balance pathogen defense with tissue protection, enhancing resistance to opportunistic microbes without exacerbating heat-induced cellular damage or algal symbiont loss^71,72^. This balance may be further fine-tuned by the temperature-sensitive regulation of steroidogenic biosynthesis pathways in *Endozoicomonas* (Fig. 5), which may exert a disproportionate influence on holobiont physiology relative to its abundance during thermal stress. Given the deep evolutionary conservation of innate immune pathways in cnidarians^73,74^, bacterially-derived steroid-based signaling may represent an underappreciated metabolic interface between bacteria and early diverging metazoa that contributes to the evolution of metazoa-microbe associations.

## Conclusion

Recent evidence suggests the origin of protosteroids occurred up to 1.6 billion years ago^75^, coinciding with the proposed evolution of unicellular eukaryotes. This correlation indicates that the origin of these molecules is most likely eukaryotic, especially given their role in sexual reproduction, which is absent in prokaryotes. Through interkingdom interactions, some bacteria might have inherited select genes from steroid hormone biosynthesis that enabled them to hijack their host immune system and thus pave the way for establishing symbiotic interactions. The limited distribution of this function across bacterial taxa associating with higher organisms suggests these microbes occupy a unique niche that distinguishes them from other microbiome members. Because of this selectivity, it may be that bacteria, such as *Endozoicomonas* that can synthesize steroid hormones, may exert a disproportionate influence on holobiont physiology compared to other bacteria. In the absence of *Endozoicomonas* during temperature-driven disease^76^, depletion of steroid precursors or reduced hormone availability may compromise host immune regulation, creating favorable conditions for opportunistic pathogens to proliferate. Our work reveals the unusual metabolic capabilities of coral-associated *Endozoicomonas* and the functional role of this bacterial genus in the coral holobiont during heat stress, as they prime the immune system of their hosts to withstand stressful events. We therefore propose a broader role for steroids in the coral holobiont and highlight the significance of understanding coral symbiosis in efforts to protect endangered coral reef habitats in a changing climate.

## Data Availability

Metagenomic reads of the coral holobiont community consortium are deposited in NCBI under the BioProject PRJNA1001615. Metagenomically assembled genomes and their annotations are available on Zenodo (https://zenodo.org/records/15276884). The untargeted mass spectrometry data is deposited on the Global Natural Products Social Molecular Network server (https://gnps.ucsd.edu/) as raw data and can be accessed under the accession numbers MSV000092488 and MSV000097641. Raw 16S rRNA gene amplicon libraries are deposited under the BioProject PRJNA1262123 in NCBI. Raw sequences and reference genome for *E. acroporae* AT381 are deposited under the accession number PRJNA1267713. Raw transcriptomic sequences of the coral are deposited under the accession number PRJNA1414119. *De novo*-assembled transcriptome, annotations, raw and normalised expression matrices are provided under the Zenodo record number 18379496.

## Acknowledgements

This work was supported by a Tamkeen award to S.A.A. (AD179) and by the NYUAD Research Institute grants ADHPG-CGSB and CG009 to the Center for Genomics and Systems Biology (CGSB) and Mubadala ACCESS, respectively. The authors acknowledge the NYU Abu Dhabi Core Technology Platform facilities for access to mass spectrometry facilities and boats for sampling. This research was carried out on the High-Performance Computing resources at New York University Abu Dhabi. Figure 1 and 6e were created using https://www.biorender.com.

## Author contributions

MAO, DA and SAA conceived the project; MAO, TDH and DA conducted the sampling; MAO, TDH, LSYC, WJ and ARH processed the samples; TDH, SA, MAO, and JBR conducted lab experiments; AM, SA, MAO, WSJ, ARH and GMR carried out data processing and analysis; AM, MAO and SAA drafted the manuscript; AM, TDH, WSJ, SAA and MAO plotted all the figures; all authors contributed to manuscript preparation; SAA and JAB acquired funding for the project.

## Additional information

This paper is accompanied by extended data and Supplementary Information.

