## Supplementary material for "Steroid hormones produced by coral-associated *Endozoicomonas* prime the coral immune response during thermal stress": ED File

### Extended Tables and Figures

**Extended Data Table 1.** Environmental metadata during coral sampling in the Arabian Gulf.

| Date | Temperature (°C) | Salinity (PSU) | DO (mg/L) | pH | Description |
|---|---|---|---|---|---|
| 28-Jun-16 | 32 | 43 | 6.2 | 8 | Early thermal stress/early disease progression |
| 29-Aug-16 | 34 | 42.2 | 5.5 | 7.9 | Severe thermal stress/further disease progression |
| 25-Oct-16 | 27.1 | 43.3 | 5.8 | 8 | Recovery |

**Extended Data Table 2.** Numbers of annotated genes from the coral host, algal symbiont and microbe assemblies; as well as predicted compounds based on *in-silico* fragmentation and putative compounds annotated based on fragmentation spectral libraries. 'Shared' indicates genes or metabolites present in two or more assemblies. Genes were annotated with eggNOG and BlastKoala, connected compounds were gathered from KEGG database.

| Assembly | Annotated genes | Predicted compounds | Putatively Annotated compounds |
|---|---|---|---|
| Host | 4619 (40.7%) | 269 (13.6%) | 43 (15.2%) |
| Algal symbiont | 620 (5.5%) | 171 (8.6%) | 42 (14.8%) |
| Microbes | 3029(26.7%) | 289 (14.6%) | 51 (18.0%) |
| Shared | 3065 (27%) | 1254 (63.2%) | 148 (52.0%) |

**Extended Data Table 3.** List of predicted biosynthetic gene clusters in MAG10 and *Endozoicomonas acroporae* AT381 genome and their predicted products based on antiSMASH.

| Cluster | Type | No. of genes | Length (bp) | Predicted molecule | Similarity score | Organism | Reference (MIBiG) |
|---|---|---|---|---|---|---|---|
| **MAG10** | | | | | | | |
| BGC01 | Ectoine | 8 | 10,395 | ectoine | 81% | *Methylophaga thalassica* | BGC0000856.1 |
| BGC02 | Betalactone | 11 | 13,681 | showdomycin | 60% | *Streptomyces showdoensis* | BGC0001778.1 |
| BGC03 | RiPP-like* | 7 | 6,297 | NA | NA | NA | NA |
| BGC04 | RiPP-like* | 11 | 9,793 | spicamycin | 42% | *Streptomyces fimbriatus* | BGC0001774.1 |
| BGC05 | Siderophore | 6 | 8,168 | aerobactin | 22%[#] | *Pantoea ananatis* | BGC0001499.1 |
| **AT381** | | | | | | | |
| BGC01 | Betalactone | 13 | 22,956 | fengycin | 13% | *Pseudoalteromonas piscicida* | BGC0001463 |
| BGC02 | Siderophore | 12 | 16,929 | aerobactin | 72% | *Pantoea ananatis* | BGC0001499.1 |

*RiPP refers to other unspecified ribosomally synthesized and post-translationally modified peptide products.

[#]Low score is based on the incompleteness of the gene cluster (loss of TonB-receptor).

**Extended Data Table 4**. Presence and absence of gene clusters inferred by pangenome related to aerobactin biosynthesis/transport (*iucC, iutA*) and degradation of steroid hormones (*kstD*), galactose (*galM*), and histidine (*hutH*). Genomes are colored based on host type (light blue, hard coral; dark blue; soft coral; orange, sponge; grey, other invertebrates; red, free-living).

| Genome | *iucC* | *proximal iutA* | *distal iutA* | *kstD* | *galM* | *hutH* |
|---|---|---|---|---|---|---|
| *Endozoicomonas* MAG10 | + | - | + | + | + | + |
| *Endozoicomonas acroporae* AT381 | + | - | + | + | + | + |
| *Endozoicomonas acroporae* Acr14 | + | - | + | + | + | + |
| *Endozoicomonas acroporae* Acr1 | + | - | + | + | + | + |
| *Endozoicomonas acroporae* Acr5 | + | - | + | + | + | + |
| *Endozoicomonas* sp. ISHI1 | + | - | + | - | - | + |
| *Endozoicomonas* sp. ONNA1 | + | - | + | - | - | + |
| *Endozoicomonas* sp. YOMI1 | + | - | + | - | - | + |
| *Endozoicomonas* sp. G2 | - | - | + | - | - | + |
| *Endozoicomonas* sp. 4G | - | - | + | - | - | + |
| *Endozoicomonas* sp. ONNA2 | - | - | + | - | + | + |
| *Endozoicomonas* sp. SESOKO1 | - | - | + | + | + | + |
| *Endozoicomonas marisrubri* | + | - | + | - | - | + |
| *Endozoicomonas* W5_Kt MAG | - | - | + | + | + | + |
| *Endozoicomonas* HY_Ok MAG | - | - | + | + | + | + |
| *Endozoicomonas* plutMAG | - | - | + | - | + | - |
| *Endozoicomonas* sp. SCSIOW0465 | - | - | + | + | + | + |
| *Endozoicomonas montiporae* LMG24815 | - | - | + | - | + | + |
| *Endozoicomonas montiporae* CL33 | - | - | + | - | + | + |
| *Endozoicomonas euniceicola* | + | - | + | + | + | + |
| *Endozoicomonas gorgoniicola* | + | - | + | + | + | + |
| *Endozoicomonas* sp. OPT23 | - | - | + | + | - | + |
| *Endozoicomonas numazuensis* | - | - | + | - | - | + |
| *Endozoicomonas arenosclerae ab112* | + | - | + | - | - | + |
| *Endozoicomonas* sp. Mp262 | - | - | + | - | + | + |
| *Endozoicomonas atrinae* | - | - | + | + | + | + |
| *Endozoicomonas elysicola* | - | - | + | - | + | + |
| *Endozoicomonas ascidiicola* | + | - | + | + | + | - |
| *S. marinus* SM1973 | + | + | - | + | - | + |

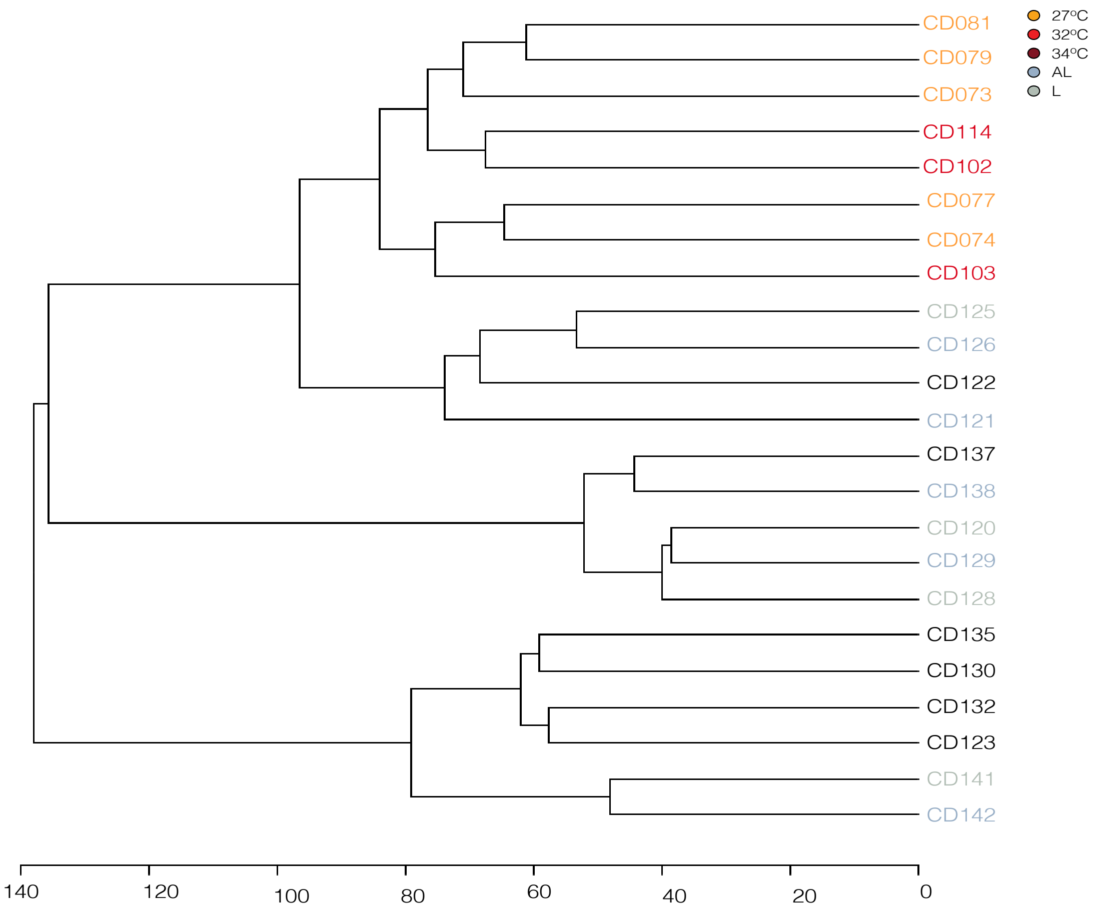


**Extended Data Figure 1. Metabolomic composition of the *A. pharaonis* holobiont across samples.** Hierarchical clustering of the holobiont metabolomes based on 5878 putative molecular features. All features were mean-centered, divided by the range of each variable, and clustered by average Euclidean distance. Refer to Extended Data Table 1 for metadata related to each sample.

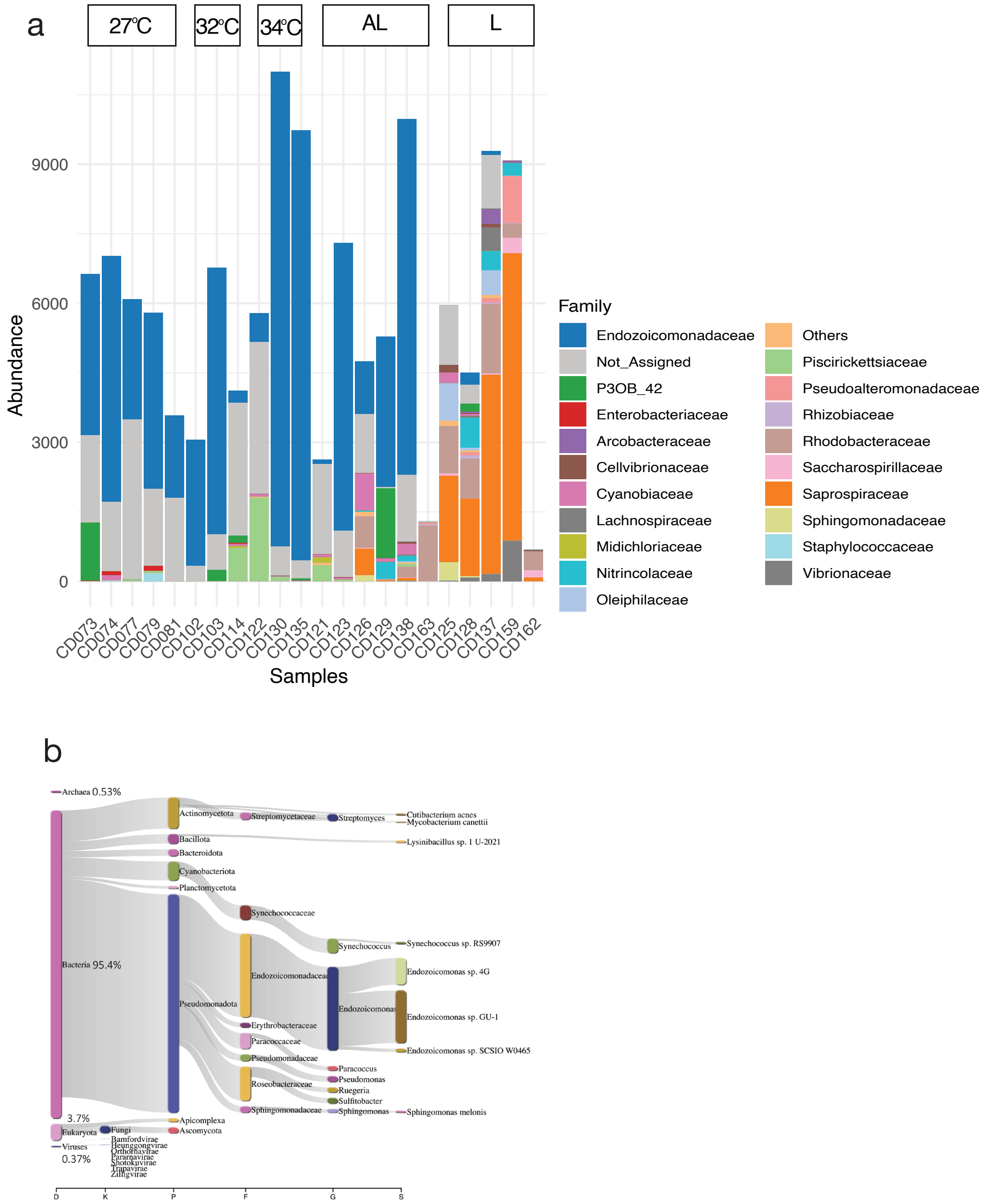


**Extended Data Figure 2. Diversity and composition of the *A. pharaonis* microbiome. a,** Taxonomic profile and abundance of amplicon sequence variants (ASVs) assigned at the family level across samples. Only the top 20 families were plotted. **b**, Read composition of putative microbial reads across *A. pharaonis* metagenomes (n = 14).

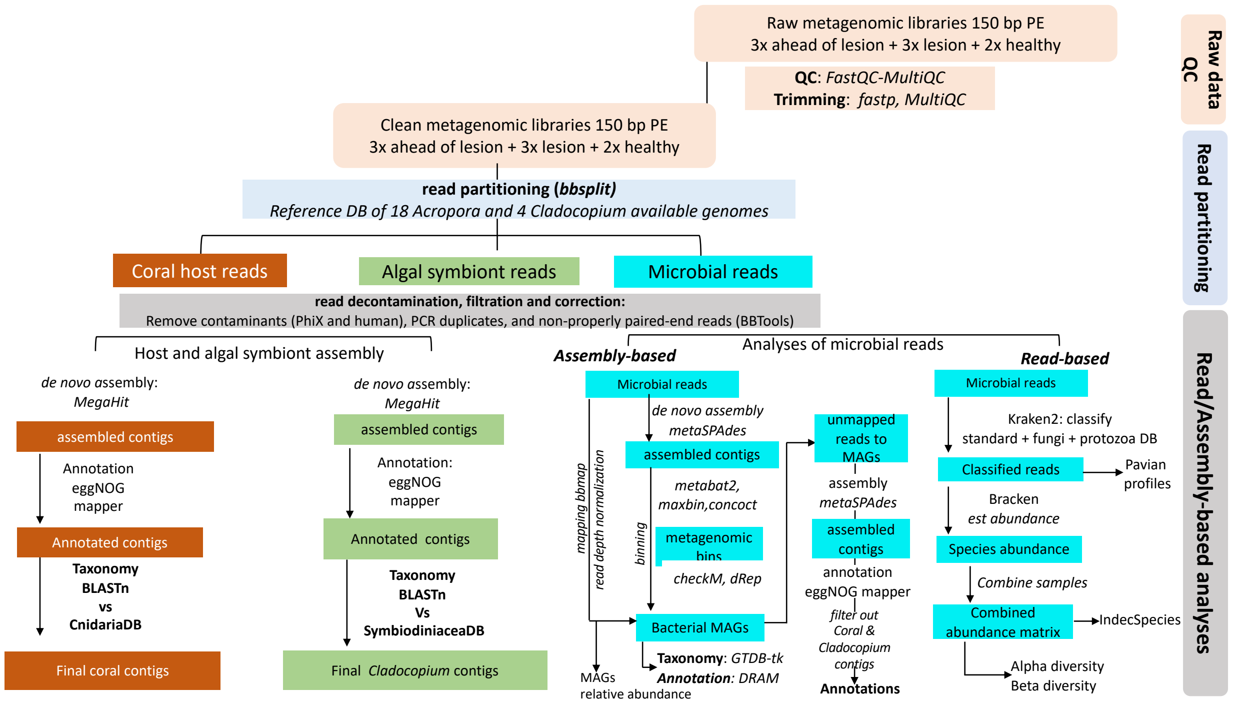


**Extended Data Figure 3. Bioinformatic workflow for metagenomics.** The pipeline used for partitioning the coral holobiont metagenomes, assembly of coral, *Cladocopium* contigs, and obtaining metagenomically-assembled genomes (MAGs) from microbes.

a

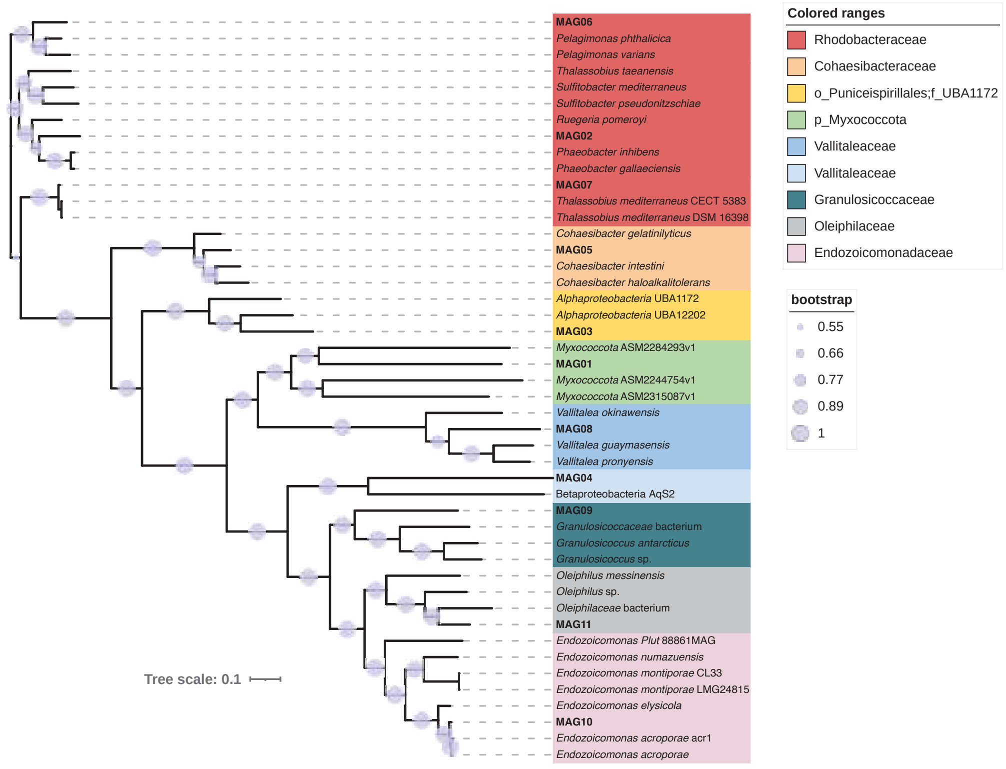


b

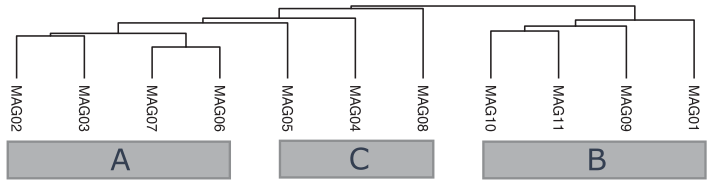


c

| KEGG_MODULE | Associated group | accession | sample_ids | Adjusted q_value |
|---|---|---|---|---|
| D-Glucuronate degradation, D-glucuronate => pyruvate + D-glyceraldehyde 3P | B | M00061 | MAG10 | 0.05 |
| Fumarate reductase, prokaryotes | B | M00150 | MAG10 | 0.05 |
| Galactose degradation, Leloir pathway, galactose => alpha-D-glucose-1P | B | M00632 | MAG10 | 0.05 |
| Glycogen biosynthesis, glucose-1P => glycogen/starch | B | M00854 | MAG10 | 0.05 |
| Guanine ribonucleotide biosynthesis IMP => GDP,GTP | B | M00050 | MAG10 | 0.05 |
| Nucleotide sugar biosynthesis, galactose => UDP-galactose | B | M00554 | MAG10 | 0.05 |
| Polyamine biosynthesis, arginine => agmatine => putrescine => spermidine | B | M00133 | MAG10 | 0.05 |
| Aerobactin biosynthesis, lysine => aerobactin | B | M00918 | MAG10 | 0.05 |
| Biotin biosynthesis, pimeloyl-ACP/CoA => biotin | B | M00123 | MAG01,MAG10,MAG11 | 0.047 |
| Cytochrome bd ubiquinol oxidase | B | M00153 | MAG10,MAG11 | 0.047 |
| Histidine degradation, histidine => N-formiminoglutamate => glutamate | B | M00045 | MAG09,MAG10 | 0.047 |

**Extended Data Figure 4. Taxonomy placement and functional enrichment analyses of metagenomically-assembled genomes (MAGs). a**, Phylogenomic tree of assembled MAGs and related reference genomes. Circles adjacent to branches represent bootstrap values (1000 bootstraps). **b**, Hierarchical clustering of MAGs based on the functional profiles inferred from KEGG mapping. **c**, Functional enrichment analyses based on the obtained clusters.

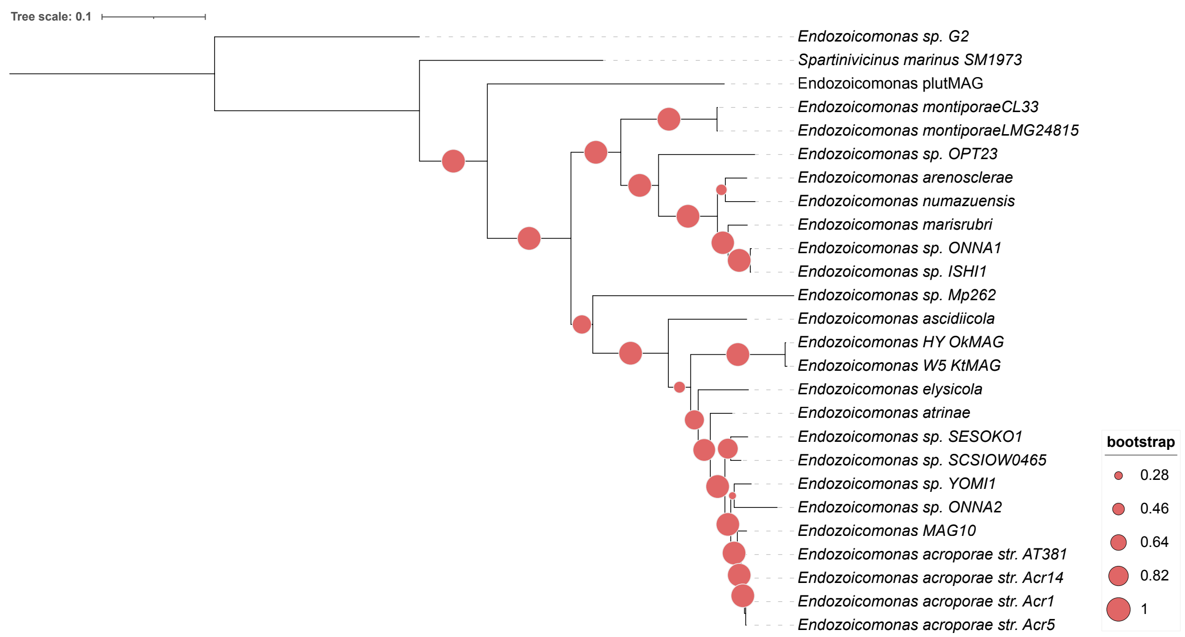


**Extended Data Figure 5. Phylogenomic tree of *Endozoicomonas* genomes.** Phylogenomic tree of *Endozoicomonas* genomes. The tree was built based on a NEWICK-formatted tree obtained from Anvio based on multiple single-copy genes identified across all genomes.

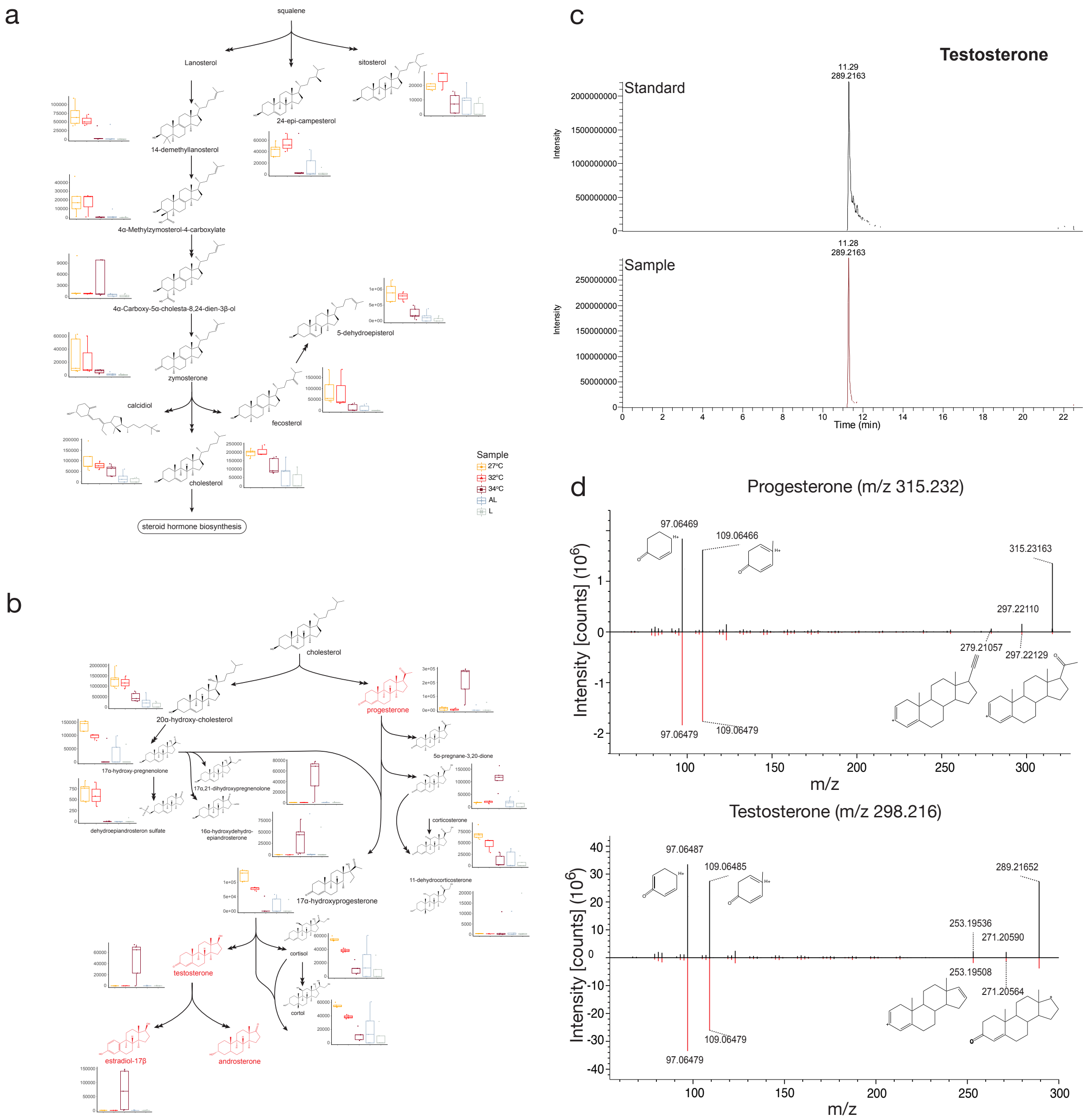


**Extended Data Figure 6. Steroid biosynthesis and steroid hormone biosynthesis and degradation (a, b) based on metagenomic and metabolomic data.** Putatively annotated metabolites detected in coral holobiont samples are shown with those highlighted in red exhibiting an unusually high relative abundance at 34°C relative to all other conditions. **c**, Retention time of testosterone captured in our samples compared to an authentic standard. **d,** MS$^2$ fragmentation spectra for progesterone and testosterone with measured spectra (black) and reference spectra (red). The structures shown indicate the characteristic fragments for each molecule.
