## Supplementary material for "Steroid hormones produced by coral-associated *Endozoicomonas* prime the coral immune response during thermal stress": SI File

**Supplementary Information Table Captions**

**Supplementary Information Figure Legends**

**Materials and Methods**

**Supplementary Information References**

### Supplementary Information Table Captions

**Supplementary Information Table 1:** List of predicted molecules by SIRIUS, their mass, retention time, and normalized abundance in samples acquired by mass spectrometry.

**Supplementary Information Table 2:** Mass fragmentation-based (see Materials and Methods) putative metabolites detected from *A. pharaonis* samples and matched with metagenome-based KEGG annotations.

**Supplementary Information Table 3:** Overview of read counts obtained from 14 coral holobiont metagenomics analysis.

**Supplementary Information Table 4:** Kegg annotation assignments across the coral holobiont metagenome

**Supplementary Information Table 5:** List of metagenomically-assembled genomes (MAGs) from this study, their GTDB taxonomy, and genome statistics.

**Supplementary Information Table 6**: List of *Endozoicomonas* genomes used for the pangenome and aerobactin biosynthesis analyses. Bold-faced genomes indicate metagenomically assembled genomes (MAGs). Only genomes with a completeness cutoff of >89% were used for isolates and >78% for MAGs. Genomes are colored based on host type (light blue, hard coral; dark blue; soft coral; orange, sponge; grey, other invertebrates; red, free-living).

**Supplementary Information Table 7:** KEGG-based Steroid biosynthesis, BGCs and metabolic functions present (+) or absent (-) in *Endozoicomonas* MAG10 compared to host and algal symbiont from metagenomic analysis.

**Supplementary Information Table 8:** MAG10 genes annotated with the Carbohydrate-Active enZYmes (CAZy) Database

**Supplementary Information Table 9:** dN/dS ratios for the siderophore receptor gene iutA and the core housekeeping gene DNA Polymerase III subunit delta across multiple *Endozoicomonas* strains. Ratios were calculated using *E. acroporae* AT381 as the reference.

**Supplementary Information Table 10:** ANOVA results examining changes in putative steroid hormones and their degradation products for the temperature gradient. P-values have been adjusted using the Benjamini-Hochberg (BH) method.

**Supplementary Information Table 11:** Mass spectrometer fragments (MS2) of steroid products produced by *E. acroporae* AT381, compared to database entries from mzCloud fragmentation database of standards.

**Supplementary Information Table 12:** T2 heat stress vs T1 control differentially expressed genes (DEGs) at FDR < 0.05.

**Supplementary Information Table 13:** T2 Progesterone + heat stress vs T1 control differentially expressed genes (DEGs) at FDR < 0.05.

**Supplementary Information Table 14:** T2 Testosterone + heat stress vs T1 control differentially expressed genes (DEGs) at FDR < 0.05.

### Supplementary Information Figure Legends

**Supplementary Information Figure 1.** $^{1}$H Nuclear Magnetic Resonance (NMR) Spectrum of androstenedione in $CDCl_3$.

**Supplementary Information Figure 2.** $^{1}$H-$^{1}$H COSY NMR spectrum of androstenedione in $CDCl_3$.

**Supplementary Information Figure 3.** $^{13}$C NMR spectrum of androstenedione in $CDCl_3$.

**Supplementary Information Figure 4.** HSQC NMR spectrum of androstenedione in $CDCl_3$.

### Materials and Methods

***Sampling.*** Samples were collected at Saadiyat Reef, Abu Dhabi UAE (N 24°35'54.2", E 54°25'12.9") by SCUBA at a depth of 6-7 m from *Acropora pharaonis* corals[1,2]. Colonies were monitored for 6 months before sampling started in June, August and October 2016. Environmental parameters (GPS coordinates, temperature, salinity, pH, dissolved oxygen, turbidity and redox potential) were measured *in situ* with a YSI Multimeter (Xylem, USA) on each sampling day (Extended Data Table 1). Nubbins of ~5 cm length were cut off with a garden shear and transferred to sterile ziplock bags filled with surrounding seawater. On the surface, nubbins were removed from the bag, placed on aluminum foil, washed with a small amount of sterile MilliQ-$H_2O$ to remove excess seawater, wrapped and flash-frozen in liquid nitrogen. The samples were stored at -80°C until further processing.

***DNA extraction.*** Using bleach-cleaned scissors and forceps, each coral fragment (~0.5 $cm^2$) was placed on dry ice, rinsed thoroughly with MilliQ-$H_2O$, gently dried and placed at the bottom of a 2.0 mL reinforced tube kept on ice. Two 3-mm stainless steel beads (acid-cleaned and sterilized) were added. Samples were incubated at 65°C for 30 min with 1 µL of RNase, Proteinase K (Promega, UK) (1 mg/ml in MilliQ-$H_2O$), lysozyme (1 mg/ml MilliQ-$H_2O$) (Sigma, Germany) and 600 µL Qiagen kit lysis buffer. Samples were homogenized twice for 10 min at medium speed with a Tissuelyzer (Qiagen, USA). Cell debris/coral skeleton was removed by centrifugation at 18,000g for 10 min in a 5430 microcentrifuge (Eppendorf, USA) at room temperature. The resulting supernatant was transferred to a fresh sterile low-bind microcentrifuge tube (Eppendorf, USA) and DNA purified using the Qiagen All-prep-Kit (Qiagen, USA) according to the manufacturer's instructions. DNA was further purified using magnetic beads (AMPure XP system, USA) with two sequential ethanol washing steps.

**16S rRNA gene amplicon sequencing.** DNA samples were prepared at 10–50 ng/µL according to Qubit 2.0 (Thermo Fisher Scientific, Waltham, MA, USA) measurements and were delivered on dry ice to the Carver Biotechnology Center (University of Illinois-Urbana Champagne, Champaign and Urbana, IL, USA) for sequencing. Briefly, libraries were prepared with the hypervariable region V4 515F (new) GTGYCAGCMGCCGCGGTAA and V4 806R (new) GGACTACNVGGGTWTCTAAT primer-pair for bacterial 16S rRNA gene amplification, with spiked PhiX as the positive control, and sequenced with MiSeq 2x250 (Illumina, CA, USA). Sequencing yielded 22 samples with sufficient reads after FASTQ check and trimming were completed. Sequences were analyzed using DADA2 v3.6.0 in R[3]. The DADA2 R-script uses raw amplicon sequencing data in FASTA format as input. Error correction of the abundances of amplicon sequence variants (ASV) was performed, quality filtered, dereplicated, chimeras removed, and end pairs were merged. For taxonomic assignment, the sequences were aligned against the SILVA database (release 132)[4]. Chloroplasts, mitochondria, and eukaryotic reads were removed. An average of 6972 reads per sample was obtained. The ASV data were then processed using the rANOMALY R package (Theil and Rifa 2021). Beta diversity was assessed and visualized through principal coordinate analysis (PCoA) based on pairwise Bray-Curtis distances and tested using permutational analysis of variance (PERMANOVA). All plots were generated using ggplot2[5] (Wickham 2016) and phyloseq (McMurdie and Holmes, 2013) R packages[1], and all statistical analyses were conducted in R version 4.1.1. Full sequencing and processing statistics can be found in the Supporting dataset Table 'Dada2 statistics'.

**Shotgun metagenomic sequencing.** DNA samples were prepared following the NovoSeq Nano DNA Sample Preparation Guide. Sequencing was performed using an Illumina Novoseq 6000 (Illumina Inc., San Diego, CA, USA) with 2x150 bp read length at Novogene (Tianjin, China). A total of 14 metagenomic libraries (3 replicates x 5 conditions) were prepared with 50 ng/mL concentration. Only one sample was not sequenced for the 27°C condition due to an inadequate amount of gDNA. Sequencing produced a total of 590 Gb of data (on average 42 Gb per sample) and, on average 280 million paired-end reads per sample (Supplementary Information Table 3).

**Quality control and *in silico* partitioning of coral holobiont metagenomes.** Illumina reads were checked for quality using fastqc v0.11.9 (https://www.bioinformatics.babraham.ac.uk/projects/fastqc/) and MultiQC v1.11[7]. Adapter sequences were trimmed, and sequences of poor quality (below Q score threshold of 30) were removed using fastp v0.23.2[8]. The *bbsplit.sh* function within the BBMap alignment tool v38.93[9] was used with default parameters in a multi-threaded process that simultaneously mapped the reads into chimeric coral and algal references obtained by concatenating the available 18 *Acropora* genomes[10] and three *Cladocopium* genomes[11–13]. This mapping strategy assigned the reads to coral, algal symbiont, or otherwise unmapped reads that potentially contained the microbial reads and hence were used for downstream analyses.

**Taxonomic classification and microbial diversity based on metagenomes.** The Kraken software suite[14] was used for read-based metagenomic analyses. Kraken2 v2.1.2[15] was used to classify the reads with the "--paired", "--report-zero-counts" and "--use-names" parameters, then BRACKEN v2.6.2[16] was used to compute species abundance by using -r 150 -l S options, and finally a species abundance matrix was obtained by combining results from all samples. The obtained species abundance raw counts matrix and taxonomy labels were used for downstream analyses. The obtained taxonomic classification data were visualized in Pavian[17]

**Pre-assembly quality control of raw metagenomic reads.** The ATLAS metagenomic pipeline v2.8.1[18] was used for read processing, assembly, and annotations. Paired reads were processed before *de novo* assembly using the BBTools suite v37.99[19] remove PCR duplicates using *clumpify*, 2) remove potential contaminants by providing the human and PhiX genomes using *BBSplit*, 3) filter reads based on quality and length using *BBduk* (trimq=10 minlength=51), and 4) correct overlapping paired-end reads based on *k*-mer coverage using *Tadpole* and merged with *bbmerge* before assembly.

**Assembly and functional annotation of coral, algal symbiont, and potential microbial contigs.** Reads that belong to the coral host or algal symbiont, based on read partitioning of the holobiont metagenome, were corrected and processed as above before *de novo* co-assembly into contigs using MEGAHIT v1.2.9[20] with default parameters. To assess the contribution of the host and algal symbiont to the functional potential of the holobiont, the assembled contigs were annotated with eggNOG mapper[21] and merged with Blastkoala/Ghostkoala[22] annotations. To ensure that the assembled contigs belonged to either coral or algal symbiont, the contigs were compared against coral and symbiont local databases using BLASTN ($e \leq 10^{-3}$). Potential microbial reads were de novo assembled individually with metaSPAdes v3.15.3[23] with default parameters. Prodigal v2.6.3[24] was used to predict open reading frames (ORFs) and the translated proteins were clustered using linclust[25] to obtain non-redundant proteins that were mapped to the eggNOG catalogue using eggNOG v5[26]. Any annotations for metazoa and dinophyta were excluded.

**Binning, annotation, and functional enrichment of metagenome-assembled genomes (MAGs).** The microbial reads were mapped to the assembled microbial contigs using *bbmap* to obtain alignment files (BAM format) required for calculating contig coverage. *metabat2* v2.15[27] uses tetra-nucleotide frequencies, differential abundance, and the presence of marker genes and was used for binning the contigs into larger bins with the "sensitive" and "min_contig_length: 1500" parameters. The obtained metagenomic bins were checked using the lineage workflow from CheckM v1.1.3[28]. MAGs were finally declared as high quality if they had completeness >75% and contamination <10%. To reduce redundancy, dRep v3.2.2[29] was used to obtain a non-redundant set of MAGs by clustering genomes assembled from all samples at a defined average nucleotide identity (ANI, 0.975) and based on genome size (>5000bp), then clusters the genomes using Mash[30] followed by MUMmer[31]. The highest-scoring bin from each cluster was only retained as the winning MAG with the highest dRep score in the dereplicated set. The relative abundance of each genome was quantified across samples based on read recruitment (mapping the reads to the non-redundant MAGs) using BBMap with default parameters. Unmapped reads to the MAGs were further assembled to contigs via metaSPAdes v3.15.3 with default parameters and annotated as previously described.

Open reading frames (ORFs) were predicted using Prodigal as described above, then Distilled and Refined Annotation of Metabolism (DRAM)[32] was used to assign database identifiers, including the Kyoto Encyclopedia of Genes and Genomes (KEGG) database, and then curate these annotations into functional categories. Genome annotations of the MAGs are provided on Zenodo (https://zenodo.org/deposit/8210635). MAGs were clustered based on KEGG modules (step coverage threshold= 0.8) using the "hclust" function implemented in R. To identify enriched functions (based on complete KEGG modules) in each group of MAGs, we used the anvi-compute-functional-enrichment that determines enrichment scores for modules within groups of MAGs by fitting a binomial generalized linear model (GLM) to the occurrence of each module in each group, and then computing a Rao test statistic, uncorrected p-values, and corrected q-values. The statistical approach for enrichment analysis is defined elsewhere[33]. We considered any function or metabolic module with a q-value equal to or less than 0.05 to be 'enriched' in its associated group.

**Taxonomic placements and phylogenomics of MAGs.** The taxonomy of the predicted MAGs was inferred using the genome taxonomy database toolkit (GTDB-Tk v5.0)[34]. GTDB taxonomy names are used throughout this paper. In addition, the GTDB-guided taxonomy of the MAGs was confirmed by phylogenomic analysis using anvio v7.1[35]. Briefly, a contig database that contains MAGs and other closely related genomes was constructed using the "anvi-gen-contigs-database" function, before hmm profiles were predicted using "anvi-run-hmms". Protein sequences of single-copy core genes were extracted and concatenated using "anvi-get-sequences-for-hmm-hits" and the phylogenomic tree was built using "anvi-gen-phylogenomic-tree". The newick-formatted tree was visualised and annotated using iTOL v5[36].

***Endozoicomonas* pangenome analyses.** *Endozoicomonas* genomes and MAGs were retrieved from the National Center for Biotechnology Information (NCBI). After filtration based on CheckM completeness cutoff of >89% for genomes and >78% for MAGs, 28 genomes (including the

*Endozoicomonas* MAG assembled in our study, MAG10) were used for the pangenome analyses following the pangenomic workflow in anvi'o v7.1[28,35]. Genome statistics, accession numbers, and host types are provided in Supplementary Information Table 4. Briefly, header lines were simplified in the 29 FASTA files using "anvi-script-reformat-fasta", fasta files were then converted to contig databases using the "anvi-gen-contigs-database". These databases were annotated using multiple functions. First, contigs were annotated with hidden Markov models (HMMs) by using "anvi-run-hmms". We then used "anvi-run-ncbi-cogs" and "anvi-run-kegg-kofams" to annotate genes in the databases with functions based on the NCBI's Clusters of Orthologous Groups (COGs) and KEGG databases. A genome storage was created by using "anvi-gen-genomes-storage" to store genomic and amino acid sequences in addition to functional annotations. Next, we used the program "anvi-pan-genome" with the default parameters to get a pangenome that includes core and accessory genomes as well as singletons. The obtained pangenome was visualized using "anvi-display-pan". The program "anvi-compute-similarity" was used to compute average nucleotide identity across genomes using the "PyANI" method. The ANI percentage identity data and Biosynthetic Gene Clusters (BGCs) predicted from AntiSMASH were added as layers into the pangenome.

**Prediction of BGCs of secondary metabolites.** Prediction of BGCs in the 29 genomes within the *Endozoicomonas* pangenome, including our assembled MAG was performed using AntiSmash bacterial v7[37] with default parameters and setting the extra features "all on". Only well-defined clusters were retained by using the "strict" option. The annotations of BGCs were assessed by matches with the Minimum Information about a Biosynthetic Gene cluster (MIBiG) database[38].

***Endozoicomonas* isolation from *A. kenti*[39], formerly *Acropora tenuis.*** Samples of three different genotypes of *Acropora kenti* (separated by 10 m) were collected from Heron Island, Great Barrier Reef, Australia, during the Summer of 2020. Fragments of *A. kenti* were rinsed with autoclaved and filtered through 0.22 µm Whatman filters(Whatman, Little Chalfont, UK) artificial seawater (AFSW), to remove any seawater-associated bacteria, and then airbrushed to obtain a tissue slurry. Coral tissue slurries of each *A. kenti* fragment were pooled and homogenized using a TissueRuptor (Qiagen) to liberate bacteria from within the tissue[39]. The pooled slurry was diluted (1/100) with AFSW, and the diluted slurry was spread on separate plates of Marine Agar 2216 (BD Difco). The agar plates were incubated at 18°C (to reduce growth of *Vibrio* spp.) until visible colonies of bacteria were observed (minimum three days). Bacterial colonies were picked and streaked onto new plates and then subsequently inoculated into 3 mL of Marine Broth 2216 (BD

Difco) at room temperature (24°C) under constant shaking (120 rpm) until maximum growth was observed. The cultured isolates were transferred into marine broth containing 20% glycerol, snap-frozen in liquid nitrogen, and stored at -80°C. The bacterial isolates were then Sanger-sequenced at the Australian Genomics Research Facility, Sydney. The resulting 16S rRNA gene sequences were matched against the NCBI database using a BLASTn search; the isolate corresponding to *Endozoicomonas* (sequence identity 96% and e-value $1e^{-67}$) can be accessed under the NCBI Sequence Read Archive (SRA) under the accession number: SAMN37396526.

**DNA extraction, genome sequencing, assembly, and annotations of *E. acroporae* AT381.** Five millilitres of overnight cultures were extracted for gDNA using the DNeasy Blood and Tissue Kit (Qiagen, Hilden, Germany) following the manufacturer's protocol. Libraries for genome sequencing were multiplexed using Nextera XT (Illumina, San Diego, US) at the Australian Genome Research Facility and sequenced on an Illumina MiSeq platform (2 × 150 bp).

Raw reads were quality filtered using fastp v0.24.0[8]. Quality-filtered and trimmed reads were then assembled into contigs in Unicycler v0.5.1[40] using SPAdes with default parameters. When assembling Illumina reads, Unicycler optimizes SPAdes by testing multiple k-mer sizes to select the best, filtering out low-depth regions to reduce contamination, applying SPAdes repeat resolution directly to the assembly graph, rejecting low-confidence repeat resolutions to minimize mis-assemblies, and trimming graph overlaps to prevent sequence repetition at contig junctions. The newly assembled genome was annotated using Bakta v1.11 (https://github.com/oschwengers/bakta), a rapid and standardized annotation tool for bacterial genomes. Genome completeness and contamination were verified using CheckM v1.2.3[28]

**Gene neighborhood analysis of aerobactin BGC *Endozoicomonas* genomes.** The coordinates of the genes within the aerobactin BGC were obtained from AntiSmash results. The operons were plotted in R using the ggplot2 v3.4.2[5] and gggenes v5[41] packages.

**Selection pressure on the siderophore receptor gene *iutA* within *Endozoicomonas*.** To investigate the selective pressures occurring on the siderophore transport gene across multiple bacterial genomes, we estimated the ratio of nonsynonymous (dN) to synonymous (dS) substitution rates (dN/dS) using the software KaKs_Calculator[42]. This software is used for unraveling the evolutionary sources acting on sequence evolution, where values of dN/dS < 1 are indicative of purifying selection: ≈1 with neutral evolution, and >1 with positive or relaxed selection. Coding sequences for the *iutA* gene and a core gene (DNA polymerase III subunit delta) were extracted from annotated assemblies using bedtools[43]. Corresponding orthologous sequences were aligned at the codon level using MUSCLE v.5[44], ensuring in-frame alignments and trimming to equal lengths as required by the KaKs_Calculator input format. We used the YN method[45] as it is robust for codon-based evolutionary analyses. dN/dS values and associated p-values were obtained for each pairwise comparison between the reference genes in *E. acroporae* AT381 and the rest.

**CAS assay.** Chrome azurol S (CAS) assay was prepared according to Schwyn and Neiland[46]. To test for Siderophore production in *Endozoicomonas*, *E. acroporae* AT381 (isolated in this study), *E. acropora* Acr14, and *E. montiporae* were ordered from the Taiwan culture collection (Bioresource Collection and Research Center, Food Industry Research and Development Institute), Taiwan. Chrome Azural S (CAS) agar plates were prepared as follows: 60.5 mg of CAS was dissolved into 50 ml of Milli Q, with the addition of 10 ml of Fe(III) solution (1 mM $FeCl_3$ in 10 mM HCl). Under slow stirring, the CAS/Fe solution was mixed with 72.0 mg Hexadecyltrimethylammonium bromide (HDTMA) dissolved into 40 ml Milli Q. Subsequently, 30 mg of PIPES buffer was dissolved into 900 ml Milli Q (pH adjusted to 6.8), with the addition of 5g of Bacto-peptone, 1 g of yeast extract and 15 g of Agar. Both solutions (Agar/PIPES and CAS/HDTMA) were autoclaved separately and later incubated at 50°C for 2 hours, and then mixed together and poured onto sterile petri dishes. *Endozoicomonas* bacterial colonies were individually picked off marine agar plates and streaked onto CAS agar. Siderophore activity was then expressed as halos forming around resultant bacterial colonies, which indicates iron acquisition by the bacteria via siderophores from CAS. In this assay, a separate negative control is not informative because strains lacking siderophore production cannot grow under iron-restricted conditions imposed by the CAS–Fe complex. Thus, absence of growth on CAS plates inherently indicates inability to extract iron.

***Metabolite extractions from coral holobionts.*** All glassware used for sample processing were baked at 420 °C for 24 hours. Coral nubbins were cut into ~1 cm sized pieces on dry ice and directly transferred into 70% methanol solution on ice in clean 20 mL scintillation vials. These were then extracted in an ultrasonication bath while cooled on ice for 30 min. The resulting slurry was transferred into 15 mL centrifuge vials (Falcon) and centrifuged at 4000 × g in a 5810R microcentrifuge (Eppendorf, USA) for 15 minutes. The supernatant was removed and transferred into pre-weighed glass test tubes, and the solvent evaporated in a Speedvac (Thermofisher, USA). Dry weight was determined, and samples were kept at -80°C until measurements.

***UHPLC-Q-ToF-MS coral holobiont metabolic profiling.*** Metabolites were resuspended in 70% methanol with 0.2% formic acid and analyzed using an Agilent 1290 HPLC system (Agilent, US) coupled to a Bruker Impact II HD Q-ToF-LC/MS (Bruker Daltonics GmbH, Germany). Metabolites were separated using a reversed-phase (RP) separation method. In RP mode, medium-polarity and non-polar metabolites were separated using an Eclipse Plus C18 column (50mm × 2.1mm ID) (Agilent, USA). Chromatographic mobile phases consisted of MilliQ-$H_2O$ + 0.2% formic acid (buffer A) and Acetonitrile + 0.2% formic acid (Buffer B). The gradient started with 95% A and 5% B, with an initial gradient of 18 min to 100% B and a holding time of 2 min. Every run was followed by a 5 min washing step cycling from buffer B to buffer A to isopropanol and equilibrated for another 2 min. Detection was carried out in positive and negative ionization modes with the following parameters: ESI settings: dry gas temperature = 220 °C, dry gas flow = 8.0 L/min, Nebulizer pressure = 2.2 bar, Capillary = 4500 V, end plate Offset = (-)500 V; MS-ToF setting: Funnel1 RF = 150 Vpp, Funnel 2 RF = 200 Vpp, hexapole RF = 50, Quadrupole = 1 eV, (Collision Energy for full scan MS = 7 eV, untargeted MSMS = stepping 30 - 50 eV); Acquisition Setting: mass range = 50–1300 m/z, Spectra rate = 6.0 Hz spectra/s, 1000 ms/spectrum. Every spectrum was individually calibrated, aligned with an internal lockmass (622.028 m/z) and peak picking was performed using the T-Rex 3D algorithm of Metaboscape v4.0 (Bruker Daltonics GmbH, Germany) with a binning threshold of <5 ppm. Background noise was removed by applying an intensity threshold of 500. Peak-picking and integration was accompanied with $^{13}C$-cluster detection for molecular feature verification.

Sirius[47], a mass fragmentation-based compound identification java tool was used to predict chemical structures and referenced against chemical databases (parameters: m/z ppm <5 ppm, Isotope score = SCORE, $MS^2$ deviation ppm < 5, All chemical formula databases, ZODIAC with default settings, CSI-Finger:ID all databases). Further, compound classes were determined using CANOPUS deep neural network fragmentation analysis[48]. Finally, the putative annotations were compared with a fragmentation library consisting of the IROA metabolite library (IROA Technologies, US). Various single metabolite standards (altogether ca. 1200 molecules) measured on the mass spectrometer and the Bruker Personal Library (15,000 Molecules- 65,000 MSMS spectra) were used to putatively annotate the metabolites with a cutoff MSMS match factor of >750 (max. 1000) and manually curated.

For statistical analysis, only features with predicted chemical structure (SIRIUS) (see above) were used in Metaboanlayst 5.0 (GenomeCanada, CAN)[49]. Data was normalized according to the internal standard chloramphenicol and range scaled (mean-centered and divided by the range of each variable)[50]. Principal component analysis (PCA) was used to estimate differences between sample groups. The statistical significance of PCA was tested using PCAtest v0.0.[51] R package.

The significance of multivariate data was tested using a linear model in Metaboanalyst 5.0[49]. Molecular features were deemed significant when achieving an adjusted p-value <0.01 (Bonferroni post-hoc tests) for FDR detection. A fold change of 1.5 was used to filter for differentially abundant compounds. Dendrograms were clustered with 'Euclidean' distance measure and 'Average' clustering algorithm. Graphs for metabolomics were performed in R (3.60) using ggplot2[5].

**Incubation of *Endozoicomonas isolates* with steroids.** Three *Endozoicomonas* strains, two isolated from *Acropora* (*E. acroporae* AT381/ Acr14) and one from *Montipora* (*E. montiporae*) were used. These strains were grown on 0.5% marine broth agar plates (35 g/L sea salts, 2.5 g/L yeast extract, 0.5 g/L protease peptone) at 34°C. An overnight culture was grown in minimal marine broth (MMB)[52]. Fifty µL of overnight culture at an optical density ($OD_{600}$) of 0.2 was used to inoculate 4 mL of sea salts (35 g/L) with 0.1% glucose as a carbon source and either cholesterol, testosterone (Cambridge Isotopes, USA), or androstenedione at a final concentration of 100 µM. All incubations were conducted with 4 replicates, except controls were done with three. The cultures were shaken at 200 rpm at 34°C for 24h. Subsequently, cultures were centrifuged at 4000 × g in a 5810R microcentrifuge (Eppendorf, USA) for 15 minutes. The supernatant was discarded, and the remaining cell pellet was directly frozen at -80°C. For extraction, the protocol mentioned above was followed.

**Measurement of metabolites in *Endozoicomonas* steroid incubations.** Liquid chromatography was performed on a ThermoFisher Vanquish UHPLC (ThermoFisher Scientific, USA) with a Kinetex C18 column (100 × 2.1 mm, 1.9 µm; ThermoFisher Scientific) under gradient conditions, using mobile phases A (0.1% formic acid in water) and B (0.1% formic acid in acetonitrile). Flow rate was set at 300 µL/min. Mobile phase A was held at 98% for 1 min followed by an increase to 100% B over 12 min and kept for 4.9 min. The gradient was then returned to 98% A within 0.1 min and re-equilibrated for 5 min. The injection volume was set at 10 µL.

Mass spectrometry analyses were performed on an Orbitrap Fusion Lumos mass spectrometer (ThermoFisher Scientific, USA) fitted with a HESI source. Source parameters were set as follows: spray voltage at + 3400 V, sheath gas at 40 (arbitrary units), auxiliary gas at 5 (arbitrary units), sweep gas at 1 (arbitrary units), ion transfer tube temperature at 300°C, and vaporizer temperature at 400°C. Ion transfer parameters were set as follows: mass range at "normal", S-lens RF level at 60%. Scan parameters applied were as follows: acquisition time from 0 to 23 min, positive ionization mode, micro scan at "1", data type at "profile", AGC target at $2 \times 10^5$. For acquisition in MS mode only, the data were acquired using the Orbitrap analyzer, at a resolution of 120,000 at *m/z* 200 (FWHM, a spectrum collected at this resolution requires ~0.2 s); maximum injection time was set at 400 ms, and the scan range from *m/z* 70 to *m/z* 700. Mass fragmentation was performed using stepped collision at 20, 30, 50 V and a resolution of 60,000 and 120 ms maximum injection time using a targeted inclusion list for predicted steroid derivatives and complemented with untargeted MS fragmentation analysis.

Raw instrument data (.RAW) were pre-processed using Compound Discoverer 3.3 SP1 (ThermoFisher Scientific, US). Metabolite detection was performed within 5 ppm mass tolerance, peak intensity cutoff of S/N 1.5. Metabolite annotations were performed using the acquired high-resolution MS/MS fragments from the samples against *mz*Cloud fragmentation database of standards (ThermoFisher Scientific, US). Authentic standards were used to identify testosterone peaks in the samples. Accurate $[M+H]^+$ *m/z*, retention time (min), and additional fragment ions *m/z* are shown in Supplementary Information Table 7 ($MS^2$-fragments). Statistical significance in the fold change was determined by calculating p-values using Student's *t*-Test[53] and adjusted using Benjamini–Hochberg approach to account for multiple testing[54].

**Field coral collections and incubation with labeled Testosterone**

Corals were collected from Snoopy Island, Fujairah (25°29′31.0″N, 56°21′48.0″E), with six colonies of *Acropora* sp. sampled and transported to the aquaria facilities at NYU Abu Dhabi (NYUAD). Colonies were maintained under ambient conditions (28°C) and acclimated for one week prior to fragmentation. Colonies were then fragmented into nubbins (2–3 cm) for experimental use and allowed to recover for an additional week in experimental aquaria prior to downstream assays. To assess coral uptake of steroid compounds, short-term incubation experiments were conducted using four glass beakers containing 150 mL of 0.2 μm-filtered seawater (Millipore), placed within a temperature-controlled water bath at 28°C. Beakers were dosed with either 10 nM, 100 nM, or 1 *μ*M of 2,3,4-$^{13}C_3$-labelled testosterone (Cambridge Isotope Laboratories, USA), or left untreated as controls. Two Acropora fragments were placed into each beaker, with gentle aeration provided via air bubblers. After 24 h and 48 h of incubation, coral fragments were removed, rinsed with 0.2 μm-filtered seawater for 5 min, immediately snap-frozen in liquid nitrogen, and stored at −80°C. An identical experiment was conducted with progesterone, though we were not able to acquire isotopically labeled progesterone. Metabolites were extracted following the holobiont extraction protocol described above; the peaks of the 2,3,4-$^{13}C_3$-labelled testosterone and naturally-abundant progesterone in the extracted samples were detected, measured and confirmed using LC-MS and LC-MS/MS by matching their accurate masses (within

5 ppm), RT and MS/MS in the mass range of 250-350 Da with authentic standards as previously described[55]. The peak areas of both testosterone and progesterone were normalized to dry extract weights.

**Experimental design for heat stress and steroid exposure.** For the heat-stress experiment, five *Acropora* colonies were fragmented into a total of 45 nubbins and affixed to ceramic reef plugs using reef-safe adhesive. Following the post-fragmentation acclimation period described above, nubbins were transferred to the experimental aquaria system, which comprised three independent 200 L sump systems containing live rock and circulating local seawater. Each sump supplied two replicate 50 L experimental tanks. Salinity was maintained at 39–40 ppt, reflecting collection-site conditions, and corals were exposed to a 12:12 h light:dark cycle at ~150 μmol photons $m^{-2}$ $s^{-1}$. Prior to initiating heat stress, coral tissue samples were collected to quantify ambient concentrations of testosterone and progesterone. Targeted analyses indicated that endogenous concentrations of both steroids fell within the range of 10–100 nM, informing the experimental dosing concentrations used in subsequent treatments.

Three nubbins from each colony were assigned to one of three treatments: (i) heat stress control (no steroid addition), (ii) heat stress with testosterone, or (iii) heat stress with progesterone. Nubbins were randomly distributed among replicate tanks within each independent sump system. Maximum photosynthetic efficiency ($F_v/F_m$) was measured daily as a proxy for coral health under dark-adapted conditions prior to light onset using an Aquapen fluorometer (Photon Systems Instruments, Czech Republic). Testosterone or progesterone (HPLC grade; Sigma-Aldrich, USA) was added to treatment tanks at a final concentration of 10 nM, and corals were incubated for 3 h prior to the first sampling time point (T1) for RNA sequencing. Seawater temperature was then rapidly increased from 28°C to 33°C over a 3 h period, consistent with CBASS-style heat-stress protocols[56], after which the second sampling time point was collected (T2). Steroid compounds were re-dosed at 10 nM, and heat stress was maintained for an additional 18 h before the final sampling time point (T3). Samples were flash- frozen in liquid nitrogen and stored at -80°C. Remaining coral fragments (n = 3–5 per treatment) were maintained at 33 °C for a further four-day period across all three conditions to assess longer-term photophysiological performance. Photochemical efficiency ($F_v/F_m$) was analysed using a mixed-effects model with treatment and time as fixed effects and coral fragment as a random effect, accounting for repeated measures over time. While no significant treatment × time interaction was detected, post-hoc Šídák-corrected comparisons were used to assess treatment differences at biologically relevant time points.

**Coral transcriptome sequencing, assembly, and analysis**. Total RNA samples were extracted using the RNeasy mini kit (Qiagen, USA) and were sent for transcriptome sequencing at Novogene (Shanghai, PRC). RNA integrity and quality were assessed prior to the subsequent mRNA enrichment from the coral holobionts. Then, mRNA libraries were prepared from total RNA using the Illumina Stranded mRNA Prep kit and sequenced using the Illumina Novaseq platform with 150-bp paired-end reads (Illumina, San Diego, CA, USA). Reads are available through the National Center for Biotechnology Information Sequence Read Archive (SRA).

Sequencing produced a total of 1.6 billion reads with an average of 31 million PE reads per library, with 3 replicates per condition. Illumina raw reads were checked for quality using FastQC version v0.11.9 and multiQC v1.11 for all datasets. Adapter and low-quality reads were removed using fastp v0.23.2. Clean reads were normalised using the Trinity script "insilico_read_normalization.pl", before assembling them into *de novo* transcriptome using Trinity V2.15.1[57] with a minimum contig length of 500. The obtained transcriptome was assessed based on the mapping rate of the filtered reads and completeness based on BUSCO (Benchmarking Universal Single-Copy Orthologues) V6[58]. The obtained transcriptome was filtered by mapping against a local cnidaria database that comprises genomes from several *Acropora* spp.[59], *Platygyra daedalea*[60], *Nematostella vectensis*[61], and *Exaiptasia pallida*[62]. Contigs with BLASTn significant hits (e-value < $10^{-5}$) were retained as the coral transcriptome and were used in downstream analysis. Open reading frames were predicted using TransDeoder V5.7.1 (https://github.com/TransDecoder/TransDecoder). The obtained protein sequences were annotated using KEGG pathways with eggNOG-mapper V2.1.13[63]. We employed the Trinity utility "align_and_estimate_abundance.pl" in a two-step process. First, it aligns filtered reads against the coral transcriptome using Bowtie2 V5.4[64], and then calculates transcript abundance based on read alignments using RSEM V.1.3.3[65], utilizing default parameters suitable for paired-end data. The gene-level count matrix was used for differential expression analysis using the Trinity script "run_DE_analysis.pl" with the DESeq2 method[66]. We compared all steroid-incubated and heat stress samples to untreated control samples. Genes with a false discovery rate (FDR) threshold of 0.05 were considered significantly differentially expressed. Genes related to innate immunity (annotated with KEGG) were investigated and plotted using the R package pheatmap V1.0.13. A summary of the signaling was illustrated using Biorender and Adobe Illustrator.

**Synthesis of androstenedione.** All reactions were magnetically stirred and conducted in a flame-dried single neck round-bottomed flask under a positive pressure of argon (i.e., balloon) unless otherwise noted. All purified products were dried under high vacuum. Organic solutions and volatiles were concentrated by rotary evaporation using a water bath at 30–35°C. Thin layer chromatography was performed with SiliaPlate glass back scored plates (60Å). Visualization of the spots was achieved by UV irradiation at 245/365 nm (Analytikjena, 6W/0.16A) or by potassium permanganate staining then charring. Purification of crude products was done by flash column chromatography (Biotage Selekt) using silica (particle size 230-400 mesh) unless otherwise noted. Yields refer to spectroscopically ($^1$H, $^{13}$C) pure materials unless otherwise noted.

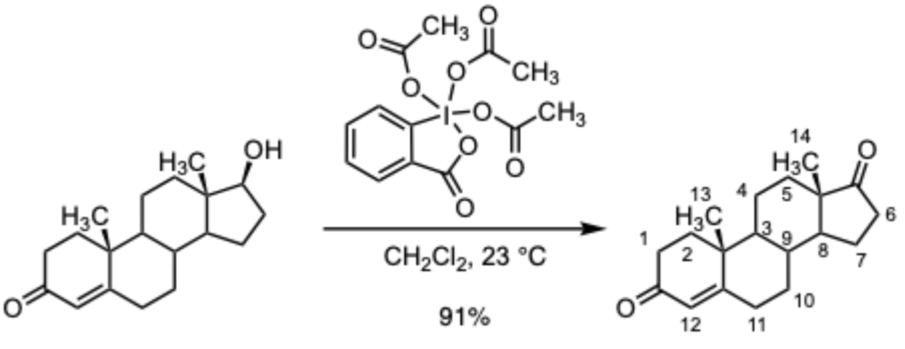


**Testosterone** **Androstenedione**

To a solution of testosterone (600 mg, 2.08 mmol, 1 equiv) in dichloromethane (105 mL, C = 0.020 M) Dess–Martin periodinane (3.53 g, 8.32 mmol, 4.00 equiv) was added. The reaction mixture was stirred for 2 h and diluted with water (150 mL). The diluted product mixture was transferred to a separatory funnel and the layers that formed were separated. The aqueous layer was further extracted with dichloromethane (2 x 50 mL), and the combined organic layers were dried over sodium sulfate. The dried solution was filtered, and the filtrate was concentrated. The residue was purified by flash column chromatography (eluting with 10% ethyl acetate–hexane initially, then a linear gradient to 30% ethyl acetate–hexane) to afford the androstenedione as a white solid (540 mg, 91%).

$R_f$ = 0.49 (50% ethyl acetate–hexane; UV)

$[\alpha]_D^{20}$: +183 (*c* 0.55, $CHCl_3$)

**$^1$H NMR** (500 MHz, $CDCl_3$) δ 5.74 (s, 1H, $H_{12}$), 2.51 – 2.25 (m, 5H, 2 × $H_1$, 2 × $H_{11}$, 1 × $H_6$), 2.16 – 2.06 (m, 1H, 1 × $H_2$), 2.06 – 2.00 (m, 1H, 1 × $H_2$), 1.99 – 1.93 (m, 2H, 1 × $H_7$, 1 × $H_{10}$), 1.89 – 1.81 (m, 1H, 1 × $H_5$), 1.77 – 1.62 (m, 3H, 1 × $H_4$, 1 × $H_6$, $H_9$), 1.55 (tt, *J* = 12.5, 9.1 Hz, 1H, 1 ×

$H_7$), 1.44 (qd, *J* = 13.2, 4.2 Hz, 1H, 1 × $H_4$), 1.34 – 1.22 (m, 2H, 1 × $H_5$, $H_8$), 1.19 (s, 3H, $H_{13}$), 1.10 (qd, *J* = 12.9, 4.3 Hz, 1H, 1 × $H_{10}$), 0.97 (ddd, *J* = 12.3, 10.7, 4.2 Hz, 1H, $H_3$), 0.90 (s, 3H, $H_{14}$).

**$^{13}$C NMR** (126 MHz, $CDCl_3$) δ 199.6 (C), 170.6 (C), 124.2 (CH), 53.9 (CH), 50.9 (CH), 47.6 ($CH_2$), 38.7 ($CH_2$), 35.8 ($CH_2$), 35.8 ($CH_2$), 35.2 (CH), 34.0 ($CH_2$), 32.7 ($CH_2$), 31.4 ($CH_2$), 30.8 ($CH_2$), 21.8 ($CH_2$), 20.4 ($CH_2$), 17.5 ($CH_3$), 13.8 ($CH_3$). One quaternary carbon was not observed. All NMR spectra are enclosed in the Supplementary Information (Supplementary Information Figures 1-4).

HRMS-CI (m/z): [M + H]+ calculated for $C_{19}H_{27}O_2$ = 287.2006; observed peak = 287.2003.

26. Huerta-Cepas, J. *et al.* eggNOG 5.0: a hierarchical, functionally and phylogenetically annotated orthology resource based on 5090 organisms and 2502 viruses. *Nucleic Acids Res.* **47**, D309–D314

(2019).
27. Kang, D. D. *et al.* MetaBAT 2: an adaptive binning algorithm for robust and efficient genome reconstruction from metagenome assemblies. *PeerJ* **7**, e7359 (2019).
28. Parks, D. H., Imelfort, M., Skennerton, C. T., Hugenholtz, P. & Tyson, G. W. CheckM: assessing the quality of microbial genomes recovered from isolates, single cells, and metagenomes. *Genome Res.* **25**, 1043–1055 (2015).
29. Olm, M. R., Brown, C. T., Brooks, B. & Banfield, J. F. dRep: a tool for fast and accurate genomic comparisons that enables improved genome recovery from metagenomes through de-replication. *ISME J.* **11**, 2864–2868 (2017).
30. Ondov, B. D. *et al.* Mash: fast genome and metagenome distance estimation using MinHash. *Genome Biol.* **17**, 132 (2016).
31. Marçais, G. *et al.* MUMmer4: A fast and versatile genome alignment system. *PLOS Comput. Biol.* **14**, e1005944 (2018).
32. Shaffer, M. *et al.* DRAM for distilling microbial metabolism to automate the curation of microbiome function. *Nucleic Acids Res.* **48**, 8883–8900 (2020).
33. Shaiber, A. *et al.* Functional and genetic markers of niche partitioning among enigmatic members of the human oral microbiome. *Genome Biol.* **21**, 292 (2020).
34. Parks, D. H. *et al.* A standardized bacterial taxonomy based on genome phylogeny substantially revises the tree of life. *Nat. Biotechnol.* **36**, 996–1004 (2018).
35. Eren, A. M. *et al.* Community-led, integrated, reproducible multi-omics with anvi'o. *Nat. Microbiol.* **6**, 3–6 (2020).
36. Letunic, I. & Bork, P. Interactive Tree Of Life (iTOL) v5: an online tool for phylogenetic tree display and annotation. *Nucleic Acids Res.* **49**, W293–W296 (2021).
37. Blin, K. *et al.* antiSMASH 7.0: new and improved predictions for detection, regulation, chemical structures and visualisation. *Nucleic Acids Res.* **51**, W46–W50 (2023).
38. Kautsar, S. A. *et al.* MIBiG 2.0: a repository for biosynthetic gene clusters of known function. *Nucleic Acids Res.* gkz882 (2019) doi:10.1093/nar/gkz882.
39. Pogoreutz, C. & R Voolstra, C. *Isolation, Culturing, and Cryopreservation of Endozoicomonas*

*(Gammaproteobacteria: Oceanospirillales: Endozoicomonadaceae) from Reef-Building Corals V1*. https://www.protocols.io/view/isolation-culturing-and-cryopreservation-of-endozo-t2aeqae (2018) doi:10.17504/protocols.io.t2aeqae.

52. Ding, J.-Y., Shiu, J.-H., Chen, W.-M., Chiang, Y.-R. & Tang, S.-L. Genomic Insight into the Host–

Endosymbiont Relationship of Endozoicomonas montiporae CL-33T with its Coral Host. *Front. Microbiol.* **7**, (2016).
