## Supplementary figures and images for "Steroid hormones produced by coral-associated *Endozoicomonas* prime the coral immune response during thermal stress"

### SI Figures

**Catalog of Nuclear Magnetic Resonance Spectra:**

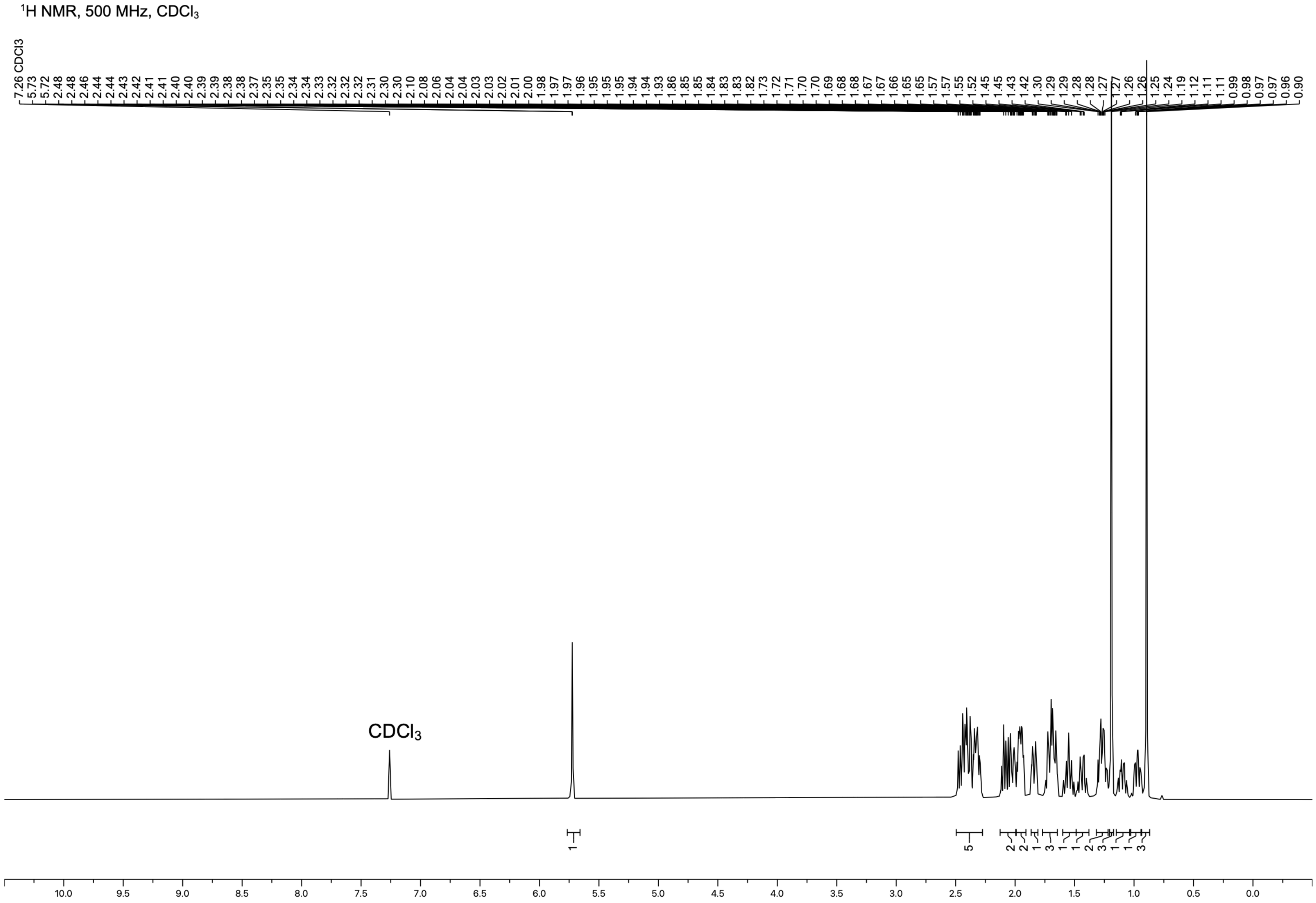

COSY, $CDCl_3$

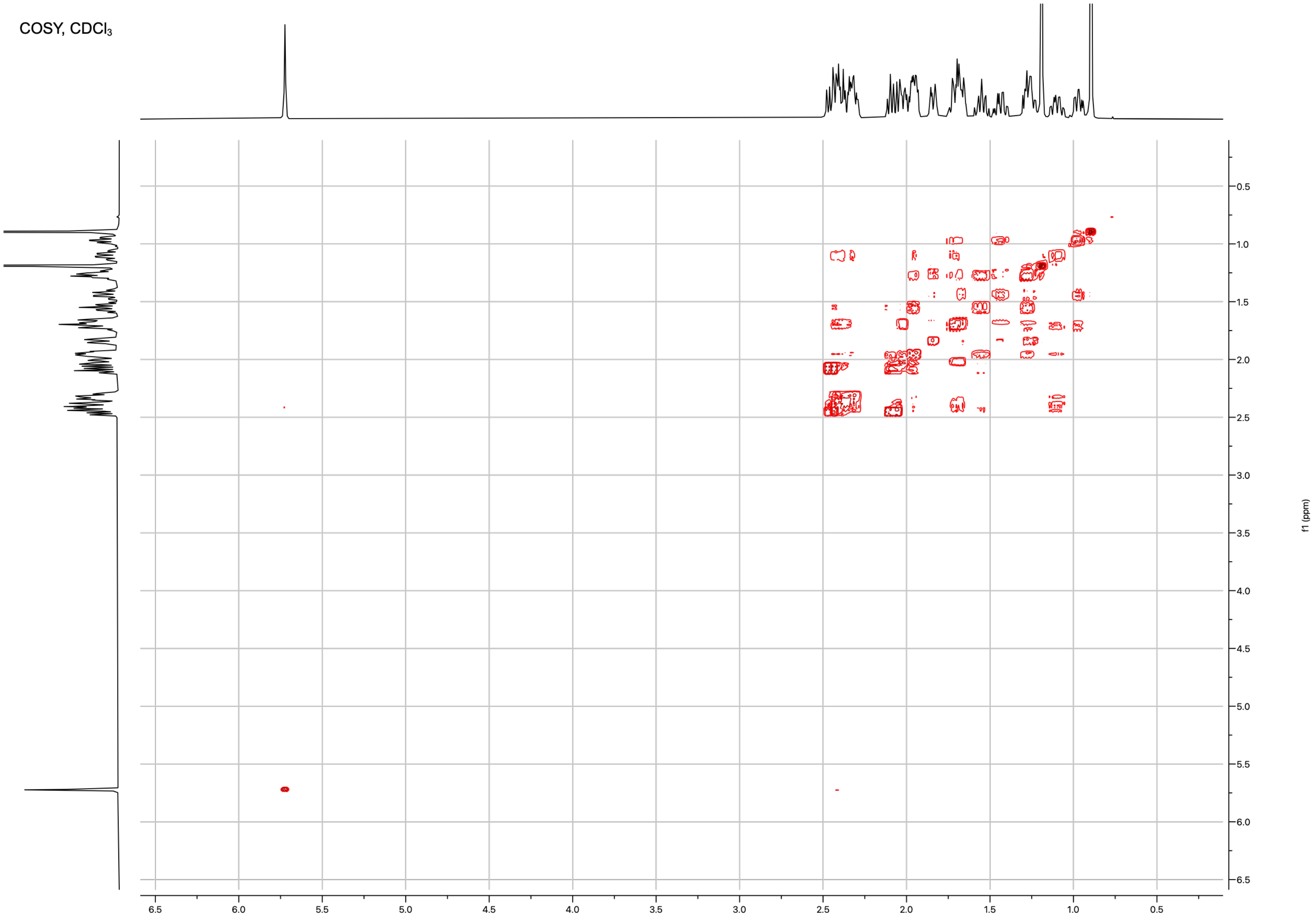

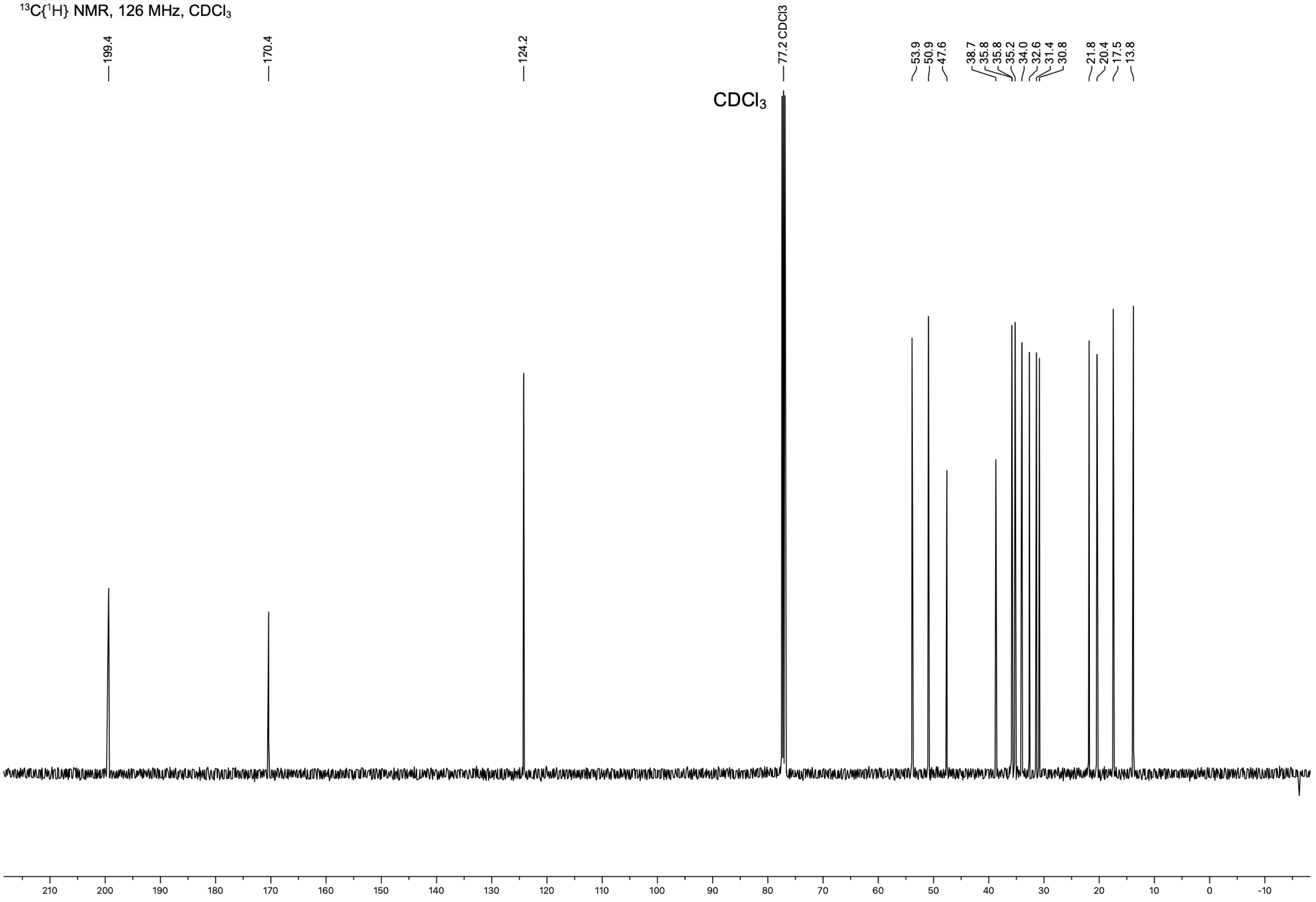

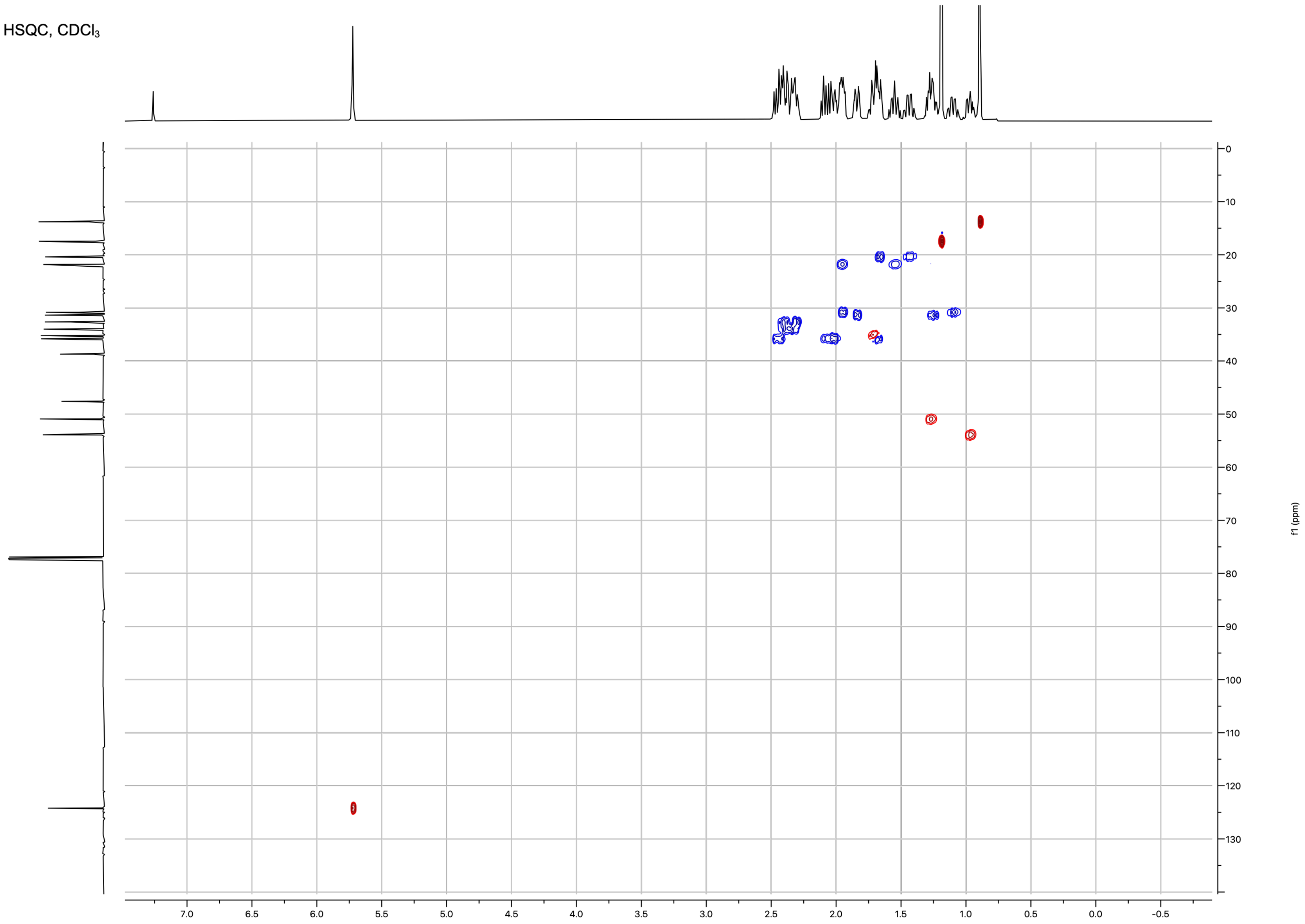
